# A mesoderm-independent role for Nodal signaling in convergence & extension gastrulation movements

**DOI:** 10.1101/671164

**Authors:** Margot L.K. Williams, Lilianna Solnica-Krezel

## Abstract

During embryogenesis, the distinct morphogenetic cell behavior programs that shape tissues are influenced both by the fate of cells and their position with respect to the embryonic axes, making embryonic patterning a prerequisite for morphogenesis. These two essential processes must therefore be coordinated in space and time to ensure proper development, but mechanisms by which patterning information is translated to the cellular machinery that drives morphogenesis remain poorly understood. Here, we address the role of Nodal morphogen signaling at the intersection of cell fate specification, patterning, and anteroposterior (AP) axis extension in zebrafish gastrulae and embryonic explants. AP axis extension is impaired in Nodal-deficient embryos, but it is unclear whether this defect is strictly secondary to their severe mesendoderm deficiencies or also results from loss of Nodal signaling *per se*. We find that convergence & extension (C&E) gastrulation movements and underlying mediolateral (ML) cell polarization are reduced in the neuroectoderm of Nodal-deficient mutants and exacerbated by simultaneous disruption of Planar Cell Polarity (PCP) signaling, demonstrating at least partially parallel functions of Nodal and PCP. ML polarity of mutant neuroectoderm cells is not fully restored upon transplantation into wild-type gastrulae, demonstrating a cell autonomous, mesoderm-independent role for Nodal in neural cell polarization. This is further demonstrated by the ability of Nodal ligands to promote neuroectoderm-driven C&E of naïve blastoderm explants in a tissue-autonomous fashion. Finally, temporal manipulation of signaling reveals that Nodal contributes to neural C&E in explants after mesoderm is specified and promotes C&E even in the absence of mesoderm. Together these results reveal a mesoderm-independent, cell-autonomous role for Nodal signaling in neural C&E that may cooperate with previously-described mesoderm-dependent mechanisms to drive AP embryonic axis extension.

## INTRODUCTION

The embryonic body plan first emerges during gastrulation, the early developmental process during which the three primordial germ layers-ectoderm, mesoderm, and endoderm- are formed and embryonic axes are manifest. The anteroposterior (AP) body axis undergoes dramatic extension at this time, a process that is essential for proper body plan formation and neural tube closure (Davidson & Keller, 1999; Wallingford & Harland, 2002). Axial extension results from highly conserved convergence & extension (C&E) movements that simultaneously elongate tissues along the AP and narrow them in the mediolateral (ML) dimension (Keller & Danilchik, 1988; Keller et al., 2000; Warga & Kimmel, 1990). This process is driven in vertebrate embryos by a combination of highly polarized cell behaviors, including mediolateral intercalation behavior (MIB) and directed migration (Keller et al., 2000; Warga & Kimmel, 1990). MIB describes the ML alignment and elongation of cells and the acquisition of bipolar protrusive behavior by which cells intercalate in a polarized fashion between their anterior and posterior neighbors (Shih & Keller, 1992a, 1992b). In vertebrate embryos, this ML polarization of cells and their behaviors is regulated by Planar Cell Polarity (PCP) signaling (Heisenberg et al., 2000; Jessen et al., 2002; Kilian et al., 2003; Park & Moon, 2002; Tada & Smith, 2000; Topczewski et al., 2001; Wallingford et al., 2000; J. Wang et al., 2006; Wang, Guo, & Nathans, 2006; Ybot-Gonzalez et al., 2007). First discovered in *Drosophila,* this conserved signaling network is required for collective polarity across cellular fields, within the plane of a tissue (Strutt & Strutt, 2005; Vinson & Adler, 1987). Core PCP components acquire asymmetric distribution within cells (Bastock et al., 2003), with some becoming enriched at the anterior or posterior aspects of vertebrate cells as they undergo gastrulation movements (Bastock, Strutt, & Strutt, 2003; Ciruna, Jenny, Lee, Mlodzik, & Schier, 2006; Roszko, S Sepich, Jessen, Chandrasekhar, & Solnica-Krezel, 2015; Y. Wang et al., 2006; Yin, Ciruna, & Solnica-Krezel, 2009). Because impairment of PCP signaling in vertebrate embryos dramatically disrupts axis extension but has little effect on patterning, it is thought to act as a molecular compass that allows cells to sense and/or respond to positional cues within the embryos (Gray, Roszko, & Solnica-Krezel, 2011; Yin et al., 2009). This implies the existence of a molecular mechanism by which patterning information is communicated to this compass, and ultimately to the cellular machinery that drives C&E cell behaviors.

In contrast with vertebrate embryos, PCP is not required for axial extension in *Drosophila,* which instead requires AP patterning conferred by the striped expression of pair-ruled genes (Irvine & Wieschaus, 1994; Zallen & Wieschaus, 2004). These in turn regulate the expression of Toll-like receptors in a partially overlapping striped pattern, thereby comprising a positional code along the extending AP axis (Paré et al., 2014). AP patterning is similarly required for extension of the gut tube in *Drosophila* and *Xenopus,* and during *Xenopus* gastrulation (Johansen, Iwaki, & Lengyel, 2003; Li et al., 2008; Ninomiya, Elinson, & Winklbauer, 2004). In particular, Ninomaya et al. (2004) reported that *Xenopus* explants with different AP positional identities extend when apposed *ex vivo,* whereas those with the same positional identity do not. Notably, these positional identities could be recapitulated by different doses of the TGFβ ligand Activin (Ninomiya et al., 2004), which signals largely via the Nodal signaling pathway (Pauklin & Vallier, 2015). These results demonstrate that patterning of the AP axis is required for axial extension, and implies a crucial role for Nodal signaling at this intersection of tissue patterning and morphogenesis.

Nodal is a TGFβ-superfamily morphogen whose graded signaling within the embryo produces discrete developmental outcomes depending on a cell’s position within that gradient and the resulting signaling level/duration to which it is exposed (Chen & Schier, 2001; Dubrulle et al., 2015; Dyson & Gurdon, 1998; Gurdon et al., 1999; van Boxtel et al., 2015). Upon binding of Nodal-Gdf3 (Vg1) heterodimers (Bisgrove, Su, & Yost, 2017; Montague & Schier, 2017; Pelliccia, Jindal, & Burdine, 2017), the receptor complex - comprised of two each of the Type I and Type II serine-threonine kinase receptors Acvr1b and Acvr2b and the co-receptor Tdgf – is activated and phosphorylates the downstream transcriptional effectors Smad2 and/or Smad3 (Gritsman et al., 1999; Schier & Shen, 2000). Nodal signaling is essential for specification of endoderm and mesoderm germ layers and patterning tissues along the AP axis, with the highest signaling levels producing endoderm and the most dorsal/anterior mesoderm fates (Dougan, Warga, Kane, Schier, & Talbot, 2003; Feldman, Dougan, Schier, & Talbot, 2000; Feldman et al., 1998; Gritsman, Talbot, & Schier, 2000; B. Thisse, Wright, & Thisse, 2000; Vincent, Dunn, Hayashi, Norris, & Robertson, 2003). Mouse embryos mutant for Nodal signaling components fail to gastrulate (Conlon et al., 1994), and Nodal-deficient zebrafish exhibit a complete lack of endoderm and severe mesoderm deficiencies (Dubrulle et al., 2015; Feldman et al., 1998; Gritsman et al., 1999), as well as neural tube closure and axis extension defects (Aquilina-Beck, Ilagan, Liu, & Liang, 2007; Gonsar et al., 2016).

Restoration of mesoderm to maternal-zygotic *one-eyed pinhead* (MZ*oep*) mutant embryos, which lack the essential Tdgf Nodal co-receptor (Gritsman et al., 1999), improves both axis length and neural tube morphology (Araya et al., 2014), strongly implying that axis extension defects in these mutants is secondary to their lack of mesoderm. Similarly, the degree of mesoderm deficiency in Nodal signaling mutants is correlated with the severity of neural tube defects (Aquilina-Beck et al., 2007; Gonsar et al., 2016). Furthermore, friction forces between anterior axial mesoderm and the overlying neuroectoderm promote axis extension (Smutny et al., 2017), again implying that Nodal regulates morphogenesis indirectly via mesoderm specification. However, the presence of mesoderm is not sufficient for C&E to occur, as demonstrated in *Xenopus* animal cap explants (Howard & Smith, 1993; Ninomiya et al., 2004). Instead, Activin signaling must be graded to drive extension (Ninomiya et al., 2004), implying an instructive rather than (or in addition to) a permissive role. Indeed, knockdown of two out of six *Xenopus* Nodal ligands disrupts C&E without affecting mesoderm formation (Luxardi, Marchal, Thomé, & Kodjabachian, 2010), implying a more direct role in morphogenesis. It is not known, however, whether or how Nodal regulates C&E of non-mesodermal tissues.

Here, we investigated the role of Nodal signaling in C&E gastrulation movements in zebrafish. We demonstrate that defective C&E movements in the neuroectoderm of MZ*oep* mutant gastrulae are associated with reduced ML cell alignment and protrusive activity. Transplantation of mutant neuroectoderm cells into wild-type (WT) gastrulae did not fully restore their ML polarity, implying a mesoderm-independent role for Nodal in cell polarization. Surprisingly, MZ*oep*-/- neuroectoderm cells exhibited normal, anteriorly-biased localization of Prickle-GFP, a hallmark of PCP polarity. Moreover, C&E in MZ*oep* mutants was reduced further by interference with the core PCP component Vangl2, demonstrating that Nodal regulates C&E cell behaviors at least partially in parallel with PCP. To further examine this cell-autonomous function of Nodal signaling in morphogenesis, we employed blastoderm explantation to isolate the effects of Nodal from endogenous signaling centers of intact embryos. We found that, as in *Xenopus* animal cap assays, expression of Nodal ligands is sufficient for robust extension of naïve zebrafish blastoderm explants in culture. Intriguingly, although these explants contain all three germ layers, their extension is driven predominantly by neuroectoderm. By combining explanted WT mesoderm with MZ*oep* mutant neuroectoderm, we find that Nodal signaling is required tissue-autonomously within the neuroectoderm for its extension *ex vivo*. Finally, treatment with Nodal inhibitors defined a late phase of signaling that promotes C&E of neuroectoderm-containing explants in the absence of mesoderm. Together, these data support a model in which Nodal signaling cooperates with PCP to promote C&E and axial extension through a combination of mesoderm-dependent and -independent mechanisms.

## RESULTS

### Nodal regulates C&E cell behaviors cell autonomously and non-autonomously

Zebrafish embryos double mutant for the two *nodal-related* genes expressed during gastrulation, *ndr1* (*sqt*) and *ndr2* (*cyc*), or lacking both maternal and zygotic function of the co-receptor Tdgf (MZ*oep*) or the downstream effector Smad2, exhibit severe mesendoderm deficiencies and impaired AP extension of the remaining neuroectoderm (Dubrulle et al., 2015; Feldman et al., 1998; Gritsman et al., 1999) (Figure 1A). However, underlying cell behavior defects during gastrulation have not been characterized. We therefore analyzed cell movements in the dorsal region of WT and MZ*oep* mutants by time-lapse confocal microscopy for a period of three hours beginning shortly after the onset of C&E movements (80% epiboly, 8.5 hours post-fertilization (hpf)). Automated tracking of fluorescently-labeled nuclei in WT gastrulae revealed clear convergence of cells from lateral positions toward the dorsal midline and concomitant extension along the AP axis (Figure 1B-C, top). Color-coding of cell tracks according to their velocities revealed that rates of cell movement are highest in the lateral-, anterior-, and posterior-most regions of the gastrula and lowest in the center (Figure 1D, top). This is consistent with mediolateral intercalation, which is characterized by a stationary point near the embryo’s equator and cell velocities that increase proportionally with their distance from this point (Concha & Adams, 1998; Glickman, Kimmel, Jones, & Adams, 2003). By contrast, MZ*oep* mutants exhibited disorganized movement and velocity patterns inconsistent with ML intercalation (Figure 1B-D, bottom). These cells moved along swirling paths rather than the direct anterior- and medial-ward movement of WT cells and were seen crossing the dorsal midline, which is not observed in WT embryos (Figure 1B)(Concha & Adams, 1998). Migration speed and track displacement were also significantly reduced in MZ*oep* mutants compared with WT gastrulae (Figure 1F), similar to results reported from slightly later developmental stages (Araya et al., 2014).

**Figure 1:**
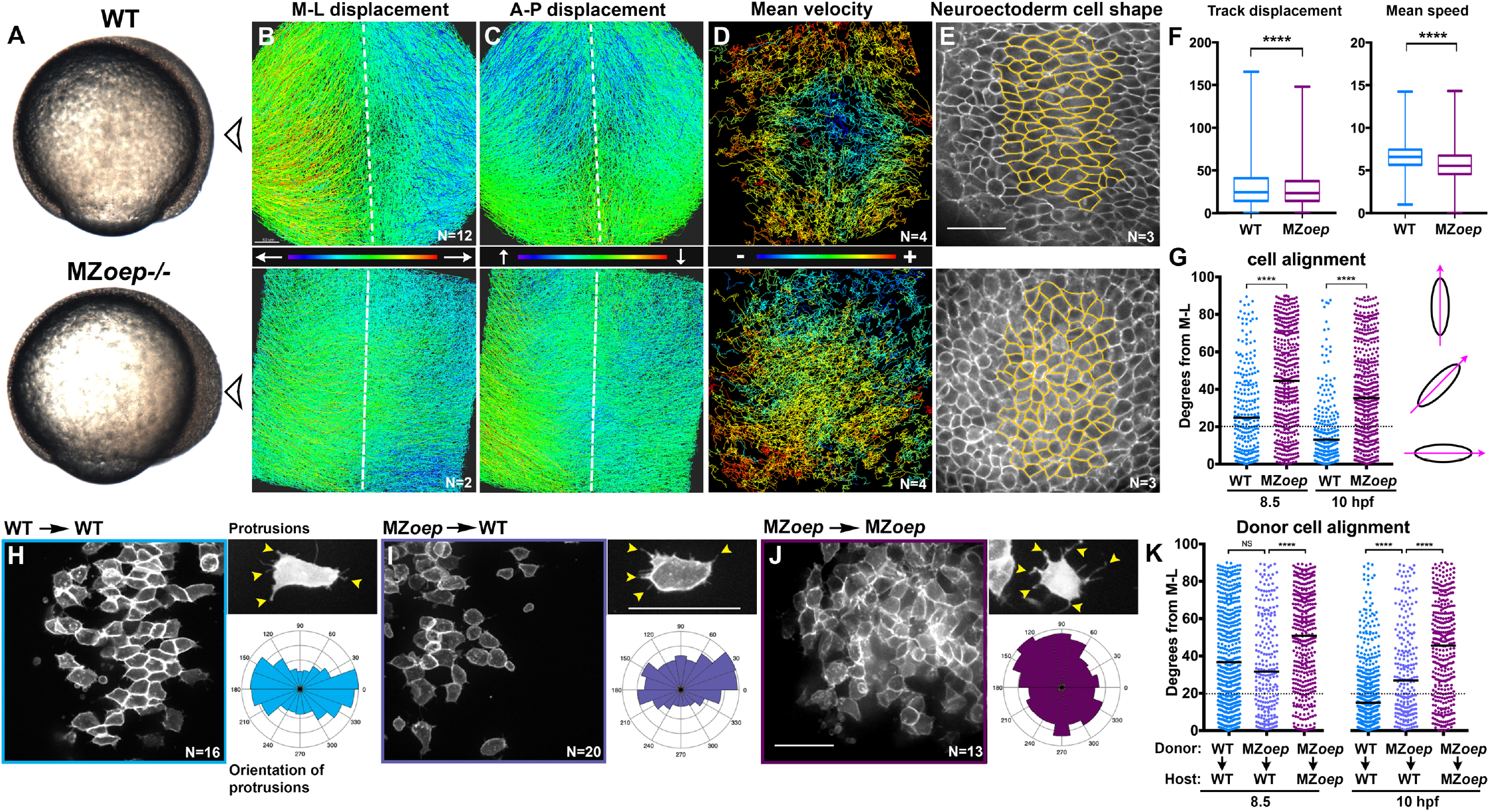
Nodal signaling regulates convergence & extension cell behaviors. **A**) Bright field images of live WT and MZ*oep*-/- embryos at 80% epiboly (8.5 hpf). Arrowhead indicates the point-of-view (dorsal side) from which all fluorescent confocal micrographs were taken. **B-D**) Representative images of automated tracking of fluorescently labeled nuclei in the dorsal hemisphere of WT (top) and MZ*oep*-/- (bottom) gastrulae. Tracks represent cell movements over three hours of time-lapse confocal imaging beginning at 8.5 hpf and are colored according to their displacement in the mediolateral (B) and anteroposterior (C) dimensions or mean velocity of cell movement (D). Dotted lines indicate dorsal midline. **E**) Representative images of membrane-labeled neuroectoderm in live WT (top) and MZ*oep*-/- (bottom) gastrulae with cells outlined in yellow. **F**) Mean speed and track displacement of labeled nuclei in WT and MZ*oep*-/- gastrulae tracked over three hours beginning at 8.5 hpf. Box plots represent the median and 25^th^ - 75^th^ quartiles, whiskers represent minimum and maximum values. N= 4 embryos of each genotype, p<0.0001, Kolmogorov-Smirnoff (K-S) tests. **G**) Neuroectoderm cell alignment at 8.5 hpf (left) and 10 hpf (right). Each dot represents a single cell, black bars are median values. N=3 embryos of each genotype, p<0.0001, K-S tests. **H-J**) Representative images of membrane-labeled donor cells within the neuroectoderm of unlabeled host gastrulae of the indicated genotypes. Right panels depict representative images (top) and orientation (bottom) of protrusions made by neuroectoderm cells of the genotypes/conditions indicated. Arrowheads indicate protrusions. Orientation of all protrusions made between 8.5 and 10 hpf is shown in radial histograms divided into 20° bins, with 0 and 180 representing ML. **K**) Donor cell alignment as in (G). The number of embryos in each condition is indicated on the corresponding panels in (H-J). Anterior is up in all images, scale bars are 50μm.

We next used a fluorescent membrane marker to assess ML cell alignment and protrusive activity underlying C&E in the neuroectoderm of WT and MZ*oep*-/- embryos from 8.5 hpf until the end of gastrulation (10 hpf) (Figure 1 E,H-J). Cell alignment was measured as the orientation of each major cell axis with respect to the embryo’s ML axis, with 0° being perfectly ML-oriented. Unlike the marked ML alignment of WT cells that increased over time (Figure 1G, blue dots), MZ*oep*-/- cells were significantly less well aligned at both 8.5 and 10 hpf time points (Figure 1G, maroon dots) (p<0.0001, Kolmogorov-Smirnov test). We then measured the orientation of cellular protrusions within the neuroectoderm of WT and MZ*oep*-/- gastrulae using cell transplantation to achieve sparse cell labeling. Protrusions made by WT cells exhibited a strong ML bias typical of MIB (Figure 1 H)(Keller et al., 2000), whereas protrusions of MZ*oep*-/- cells were essentially randomly oriented with a slight anterior bias (Figure 1J). Together, these results demonstrate a severe disruption of C&E gastrulation movements and polarized cell behaviors in MZ*oep* mutants.

Previous studies have shown that axis extension can be largely rescued by restoration of mesoderm to MZ*oep* mutant embryos (Araya et al., 2014), but the autonomy of Nodal signaling within the neuroectoderm has not been examined at the level of cell polarity. To determine whether the presence of mesoderm improves ML polarity of MZ*oep* mutant neuroectoderm cells during gastrulation, we transplanted membrane-labeled MZ*oep*-/- cells into the neuroectoderm of WT hosts (Figure 1I). We found that the degree to which polarity was restored varied by developmental stage. At 8.5 hpf, the alignment of MZ*oep*-/- cells within WT hosts was not significantly different from WT donor cells in WT hosts, whereas at 10 hpf, MZ*oep*-/- donor cells were significantly less well aligned than WT controls (Figure 1K) (p<0.0001, K-S test). Notably, mutant cells in WT host gastrulae were significantly better aligned than MZ*oep*-/- cells in MZ*oep*-/- hosts at both time points (Figure 1K) (p<0.0001, K-S test). Orientation of MZ*oep*-/- cellular protrusions was also partially improved within WT hosts, although they failed to achieve the clear bipolarity of WT protrusions (Figure 1I). These results reveal two phases of neuroectoderm cell polarization: an initial period during which Nodal-deficient cells polarize normally in a WT environment, and an additional later phase when Nodal is required cell-autonomously for full ML polarity. Together these results reveal an essential role for Nodal signaling in neuroectoderm C&E, including a discrete mesoderm-independent phase of ML cell polarization.

### Nodal functions partially in parallel with PCP signaling during axis extension

Because PCP signaling is a critical regulator of ML cell polarization underlying C&E, we next assessed the activity and polarity of this signaling network within MZ*oep* mutant neuroectoderm. We first examined expression of PCP signaling components in MZ*oep* mutants by whole mount in situ hybridization (WISH) at late gastrulation. Expression of the Wnt/PCP ligand *wnt11* is a known direct transcriptional target of Nodal signaling (Dubrulle et al., 2015; Gritsman et al., 1999), but other PCP genes *wnt5b, vangl2 (trilobite), and gpc4 (knypek)* were all expressed robustly in MZ*oep*-/- gastrulae (Figure 2A). We next transplanted cells expressing a GFP-fusion of the core PCP component Prickle (Pk-GFP) into the neuroectoderm of host gastrulae and assessed localization of Pk-GFP to the anterior edge of neuroectoderm cells as a read-out of PCP signaling activity (Ciruna et al., 2006; Yin, Kiskowski, Pouille, Farge, & Solnica-Krezel, 2008). We found that WT donor cells in WT hosts and MZ*oep*-/- donor cells in MZ*oep*-/- hosts exhibited similar proportions of anteriorly-localized Pk-GFP puncta, although the latter contained significantly more membrane-associated puncta that were not anteriorly localized (chi-square, p=0.0001) (Figure 2B-C). Finally, we tested whether disrupting PCP signaling with an antisense morpholino oligonucleotide (MO) against *vangl2* (Williams et al., 2012) further reduced C&E in MZ*oep* mutant gastrulae. Upon injection with *vangl2* MO at a dose that phenocopies *trilobite* mutant C&E defects (Figure 2G)(Solnica-Krezel et al., 1996), the neural plates of both WT and MZ*oep* mutant embryos were significantly wider and shorter than in uninjected controls at late gastrulation (Figure 2D-F), indicating further C&E reduction. Because disrupted PCP signaling exacerbated C&E defects in embryos completely devoid of Nodal signaling, together with our data suggesting largely intact PCP signaling in MZ*oep* mutants, these results indicate that Nodal functions at least partially in parallel with PCP during C&E gastrulation movements.

**Figure 2:**
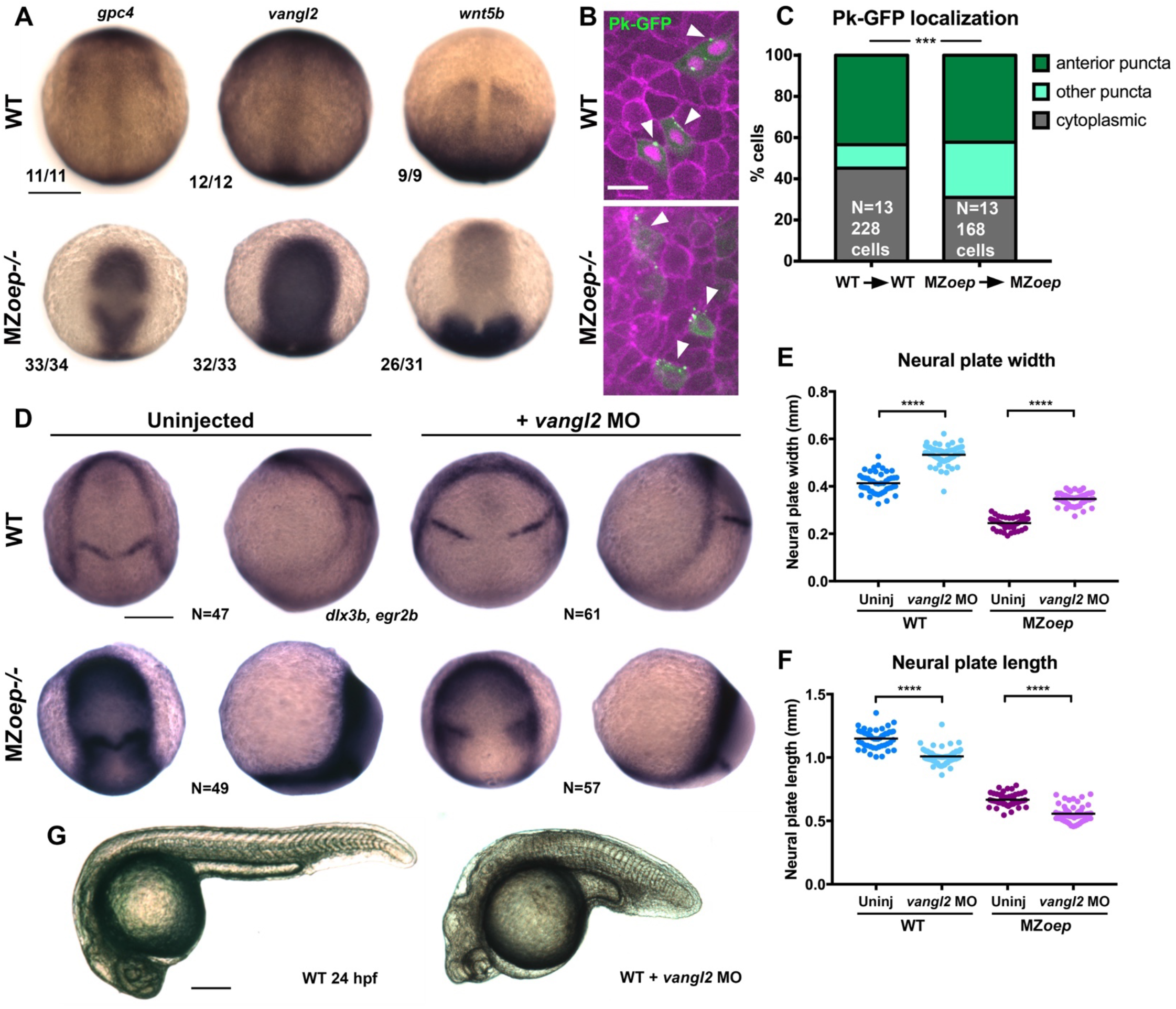
Nodal regulates C&E at least partially in parallel with PCP signaling. **A**) In situ hybridization for the transcripts indicated in WT (top) and MZ*oep*-/- (bottom) gastrulae at 9.5 hpf. Fractions indicate the number of embryos with the pictured phenotype over the number of embryos examined. **B**) Representative images of transplanted Prickle (Pk)-GFP donor cells within the neuroectoderm of membrane-labeled WT and MZ*oep*-/- host gastrulae. Arrowheads indicate puncta at anterior edges. **C**) Pk-GFP localization in the genotypes indicated. N indicates the number of embryos and cells analyzed for each condition, p<0.001, Chi-square test. **D**) In situ hybridization for *dlx3b* and *egr2b* in WT (top) and MZ*oep*-/- (bottom) gastrulae at 9.5 hpf, uninjected or injected with 2ng MO4-vangl2. Dorsal views on the left, lateral views on the right. **E-F**) Width (E) and length (F) of neural plates in the embryos depicted in (D). Each dot represents a single embryo, black bars are mean values. Number of embryos in each condition is indicated on the corresponding panel in (D), p<0.0001, Unpaired T-tests. **G**) Representative images of live WT embryos uninjected (left) or injected with 2ng MO4-*vangl2* (right) at 24 hpf. Anterior is up in (A-F), to the left in (G). Scale bars are 20μm in (B), 200μm in all other panels.

### Nodal signaling promotes ex vivo extension and tissue patterning

The results described above demonstrate that Nodal signaling is necessary for full polarization of cells and cell behaviors underlying C&E during gastrulation. To test whether Nodal is also sufficient for these behaviors, we sought to define its role during axis extension in relative isolation, independently of other signaling and patterning events within the embryo. To this end, we employed blastoderm explantation, a technique by which the animal region of blastula-stage zebrafish embryos is isolated from endogenous signaling centers at the embryonic margin to produce clusters of relatively naïve cells that can be grown and manipulated in culture (Sagerström, Grinblat, & Sive, 1996; Xu, Houssin, Ferri-Lagneau, Thisse, & Thisse, 2014). To determine the effect of Nodal signaling on such explants, we injected WT embryos at one-cell stage with a series of doses (from 2.5 to 100pg per embryo) of synthetic *ndr2* mRNA, explanted the animal portion of the blastoderm at 256-512 cell stage (2.5 hpf), and cultured them *ex vivo* until intact siblings reached two to four somite (2-4S) stage (Figure 3A, Figure S3). Several of these doses induced robust extension of explants in culture, while explants cut from GFP-injected or uninjected WT control or *ndr2*-injected MZ*oep*-/- embryos failed to extend (Figure 3B-E, Figure S3).

**Figure 3:**
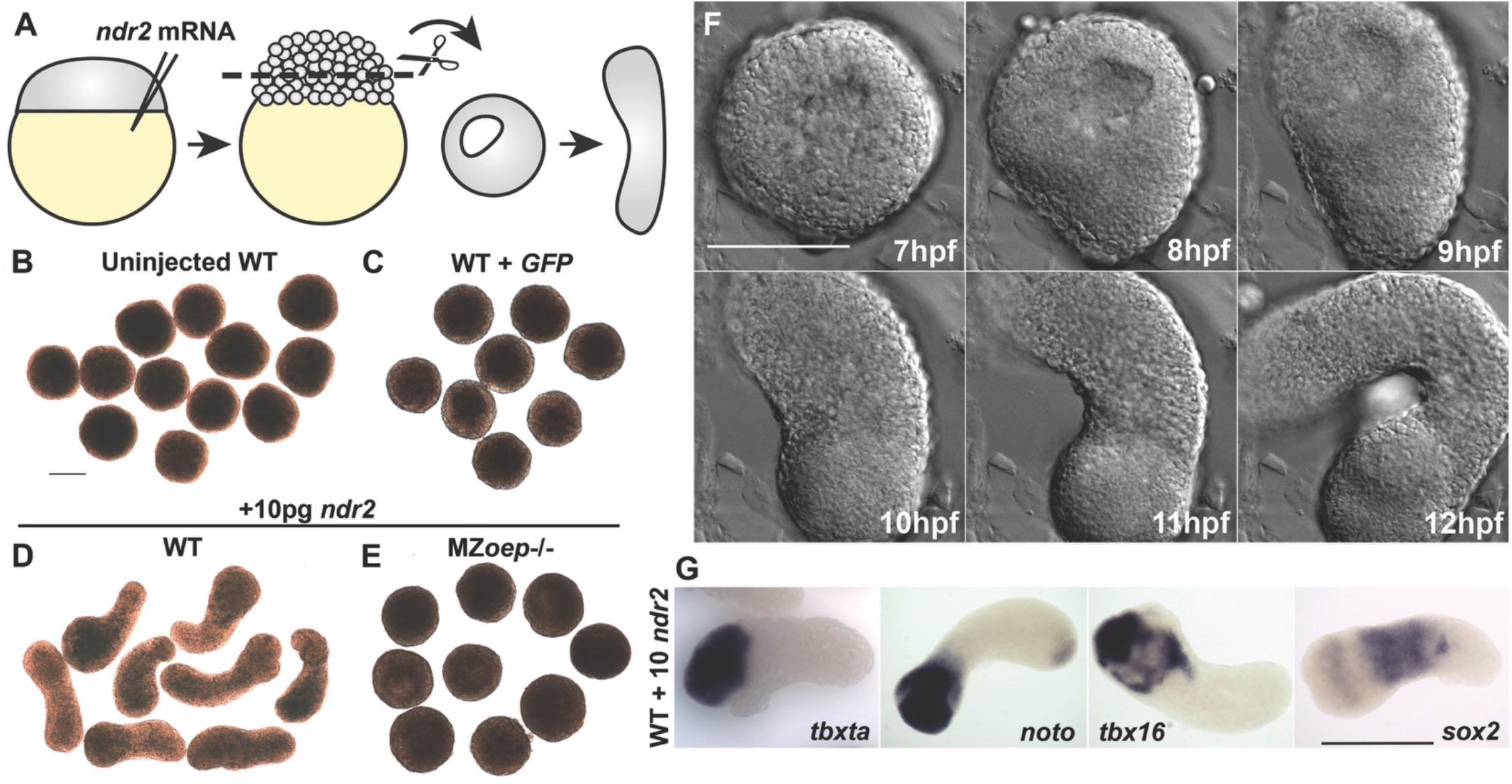
Nodal ligands promote *ex vivo* C&E of blastoderm explants. **A**) Diagram of injection and explantation of zebrafish embryos. **B-E**) Representative bright field images of live blastoderm explants of the indicated conditions/genotypes at the equivalent of two to four somite stage. **F**) Time-lapse DIC series of a WT explant injected with 10pg *ndr2* RNA. **G**) In situ hybridization for the transcripts indicated in WT explants injected with 10pg *ndr2* RNA. Scale bars are 200μm.

We assessed explant extension by live time-lapse imaging and length/width ratio measurements of fixed explants at the equivalent of 2-4S stage. Time-lapse imaging of live *ndr2*-injected explants revealed the onset of extension morphogenesis at or around 8 hpf (Figure 3F), corresponding precisely with the start of C&E movements in intact embryos. While all *ndr2* doses induced some degree of extension over control explants, the intermediate doses (5-25pg) were most effective, with 10pg producing the most extension and the highest dose, 100pg, producing the least (Figure S3B). Because it produced the most robust extension, 10pg *ndr2* was used for most subsequent experiments.

Whole mount *in situ* hybridization (WISH) revealed that explants expressed markers of endoderm, mesoderm (dorsal gastrula organizer, axial and paraxial), and neuroectoderm according to the dose of *ndr2* with which they were injected, consistent with Nodal-dependent tissue induction in intact embryos (Chen & Schier, 2001; Feldman et al., 2000; B. Thisse et al., 2000). For example, low doses of *ndr2* induced robust expression of the neuroectoderm marker *sox2* and some paraxial mesoderm *(tbx16),* whereas high doses induced expression of the endoderm marker *sox17* and the organizer gene *gsc,* but less neuroectoderm (Figure S3A). Uninjected control explants expressed none of these tissue-specific markers at appreciable levels. Notably, explants injected with 10pg *ndr2* exhibited discrete gene expression domains: mesoderm markers *tbxta, tbx16,* and *noto* were nearly always restricted to one end, *sox17* was present in small spots (likely individual endoderm cells), and *sox2* was observed in a striped pattern orthogonal to the long axis of each explant (Figure 3G, Figure S3A). Together these results demonstrate that Nodal signaling specifies a number of tissue types in discrete, spatially-organized domains and promotes C&E morphogenesis within isolated naïve blastoderm in a dose-dependent manner.

### Blastoderm explants exhibit asymmetric Nodal signaling

The gene expression patterns observed in *ndr2*-expressing explants revealed asymmetry along the axis of extension, which is known to be critical for C&E morphogenesis of *Xenopus* explants (Ninomiya et al., 2004). To test whether Nodal signaling activity could account for this asymmetry, we immuno-stained *ndr2*-injected and uninjected WT control explants for phosphorylated Smad2, an indicator of active Nodal signaling (Figure 4A). We identified pSmad2+ and pSmad2-nuclei within explants using automated spot detection and colocalization analysis (Figure 4A, see methods), then quantified their spatial distribution along the “axis” of each explant (Figure 4B-C). Comparing the distribution of Smad2+ nuclei (Figure 4B-C, blue dots) to all nuclei (gray dots) within explants revealed significant asymmetry beginning at 6 hpf and increasing through 8 hpf in *ndr2*-injected explants (p<0.0001, Mann-Whitney test), but very few pSmad2+ nuclei that exhibited little to no asymmetric distribution in uninjected controls (Figure 4B-C). Notably, the timing of pSmad2 detection differed between *ndr2*-injected explants and intact WT embryos, where an appreciable pSmad2 signal was first seen around 5 hpf (Figure S4A). pSmad2 then persisted in explants until at least 8 hpf, whereas no signal was detected after 6 hpf in embryos (Figure S4A). No signal was detectable in embryos treated with the small molecule Nodal inhibitor SB505124 (DaCosta Byfield, Major, Laping, & Roberts, 2004)(Figure S4B-C), indicating that this antibody specifically detected Nodal signaling activity. Further evidence of asymmetric Nodal signaling within explants was provided by increasing levels and asymmetry of *lefty1* (*lft1*) expression, a negative feedback inhibitor and direct transcriptional target of Nodal signaling (Meno et al., 1999), in *ndr2*- expressing but not control explants (Figure 4D). These results demonstrate that injection of *ndr2* mRNA at one-cell stage produces extending explants with asymmetric Nodal signaling activity.

**Figure 4:**
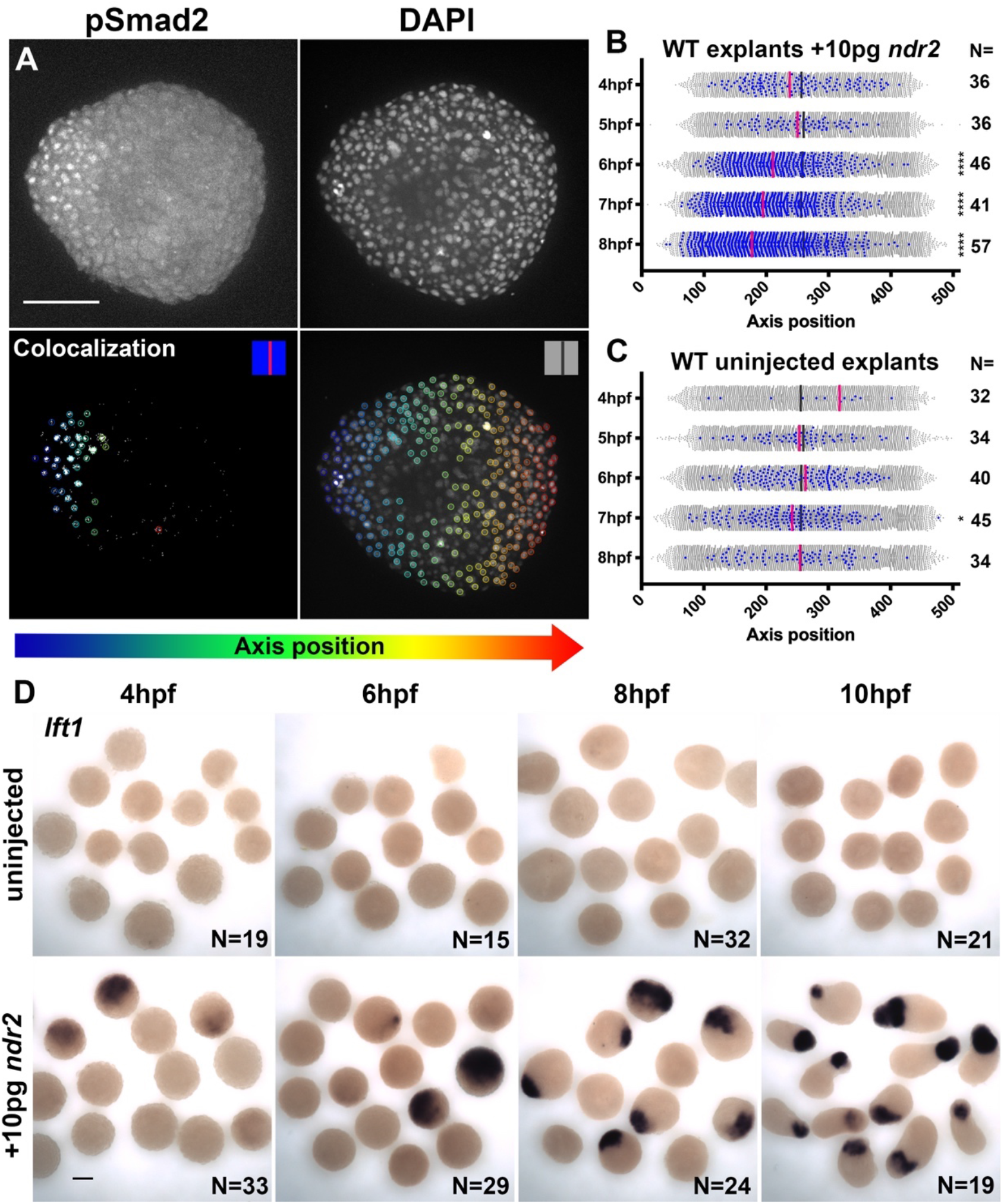
Nodal-expressing explants exhibit asymmetric Nodal signaling activity. **A**) Representative confocal z-projections and colocalization of immunofluorescent staining for phosphorylated Smad2 and DAPI-labeled nuclei in 8 hpf explants. Colored circles indicate automatically detected spots and are colored according to position along the axis of each explant. **B-C**) Axis position of pSmad2-positive nuclei (blue) and all nuclei (gray) in explants injected with 10pg *ndr2* (B) or uninjected (C) at the time points indicated. Each dot represents a single nucleus, pink and dark gray bars are median values. N= number of explants in each condition. Mann-Whitney tests to compare distribution of pSmad+ nuclei to all nuclei, ****p<0.0001, *p<0.05. **D**) In situ hybridization for *lefty1* in uninjected (top) and *ndr2*-injected (bottom) explants at the time points indicated. Scale bars are 100μm.

### Nodal signaling and PCP promote cell polarity underlying C&E ex vivo

*Xenopus* animal cap explants exposed to asymmetric Activin signaling exhibit robust ML cell polarization and intercalation (Ninomiya et al., 2004). To analyze cell polarity underlying Nodal-driven C&E in our explants, we utilized fluorescent membrane labeling to quantify cell alignment in live *ndr2*-expressing explants (Figure 5A-C). In *GFP*-expressing control explants, cells were randomly oriented (Figure 5A,C median angle=44°) when intact sibling embryos reached 2-4S stage. This was in stark contrast to *ndr2*-expressing explants, whose extension was accompanied by robust ML alignment of cells, defined as orthogonal to the axis of extension (Figure 5A,C median angle=19°), demonstrating that Nodal signaling is sufficient to induce ML cell alignment underlying C&E morphogenesis in populations of otherwise naïve cells.

**Figure 5:**
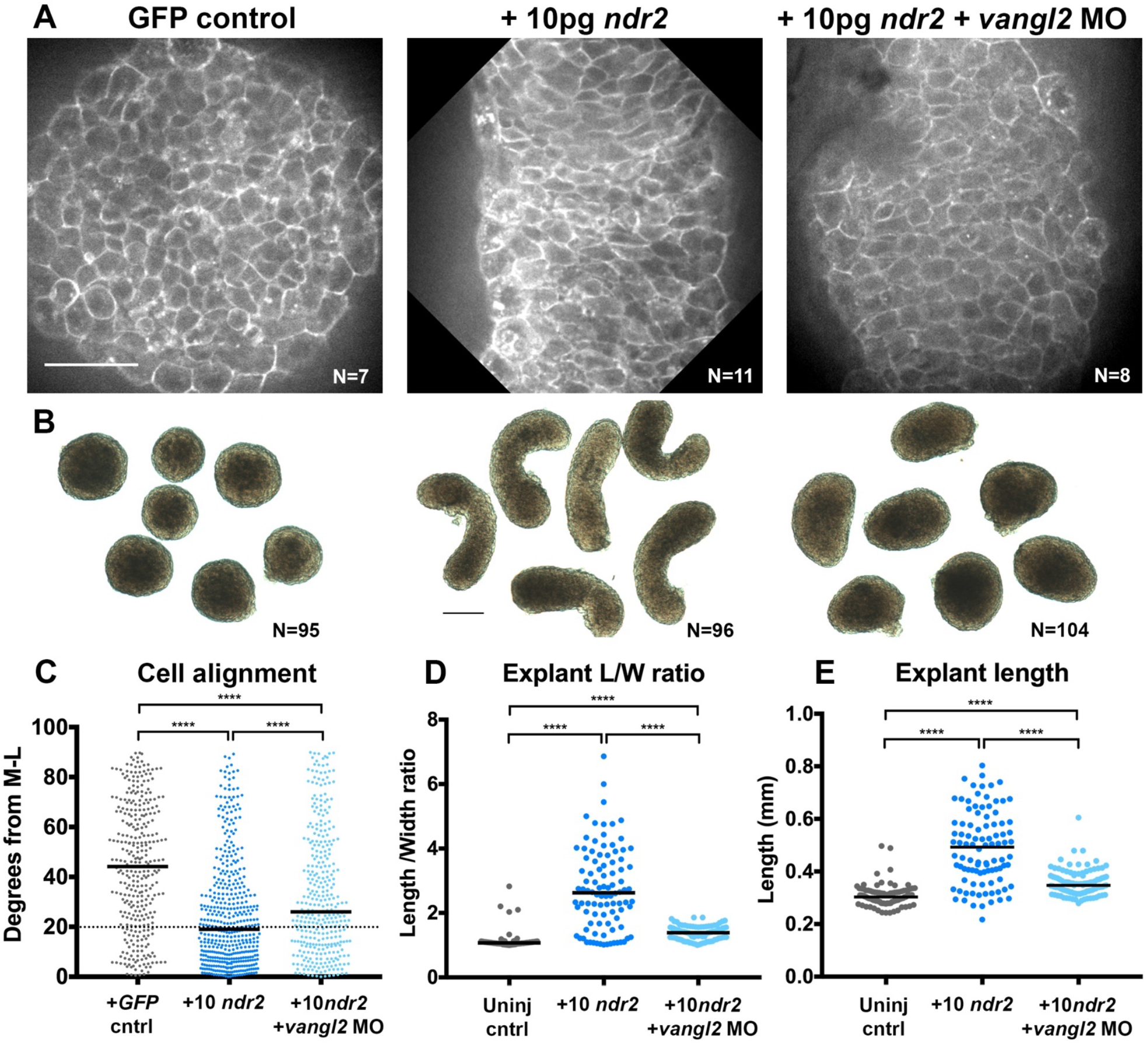
Disrupted PCP reduces Nodal-induced cell polarization and C&E within explants. **A**) Representative confocal micrographs of live membrane-labeled explants of the indicated conditions/genotypes at equivalent of 2-4 somite stage. **B**) Representative bright field images of live blastoderm explants at 2-4 somite stage. **C**) Explant cell alignment as in Figure 1. Number of explants in each condition is indicated on the corresponding panel in (A). Mediolateral (M-L) is defined as orthogonal to the axis of extension. **D-E**) Length/width ratios (D) and length (E) of explants depicted in (B). Each dot represents a single explant, black bars are median values. Number of explants in each condition is indicated on the corresponding panel in (B), p<0.0001 Kruskal-Wallis test. Scale bar is 50μm in (A), 200μm in (B).

To ask whether Nodal-dependent *ex vivo* extension and ML cell alignment require PCP signaling, we generated explants from embryos co-injected with 10pg *ndr2* mRNA and *vangl2* MO. Vangl2-deficient explants exhibited overall length and length/width ratios that were significantly reduced compared with those expressing *ndr2* alone, but significantly higher than uninjected controls (Figure 5B,D-E). Live imaging of fluorescently-labeled cell membranes further revealed that ML cell alignment was reduced, but not entirely randomized in *vangl2* morphant explants compared with those expressing *ndr2* alone (Figure 5A,C median angle=26°). Because Nodal is necessary and sufficient for ML cell polarization *ex vivo,* and this polarity is reduced upon disruption of PCP signaling, these results indicate that PCP signaling functions downstream of Nodal in explant extension. However, because *vangl2* MO reduces but does not completely abolish Nodal-induced C&E and ML cell alignment, our data are consistent with an additional PCP-independent role for Nodal signaling in C&E.

### Nodal signaling promotes ex vivo C&E of neuroectoderm

The localization of mesoderm tissue to one end of each *ndr2*-expressing explant (Figure 3) suggests that their extension is driven largely by non-mesodermal tissues. To directly observe mesoderm within explants as they extend, we performed live time-lapse confocal imaging of explants from *ndr2*-injected transgenic *lhx1a:gfp* embryos in which much of the axial and lateral mesoderm expresses GFP (Swanhart et al., 2010). Consistent with WISH in fixed explants (Figure 3), ~77% of live explants contained a mass of GFP+ cells at one end that was displaced as the GFP-region underwent extension (Figure 6A-B). This non-mesodermal portion of each explant expressed the neuroectoderm marker *sox2* by WISH (Figure 3G, Figure S3A), strongly implying that extension is driven by neuroectoderm.

**Figure 6:**
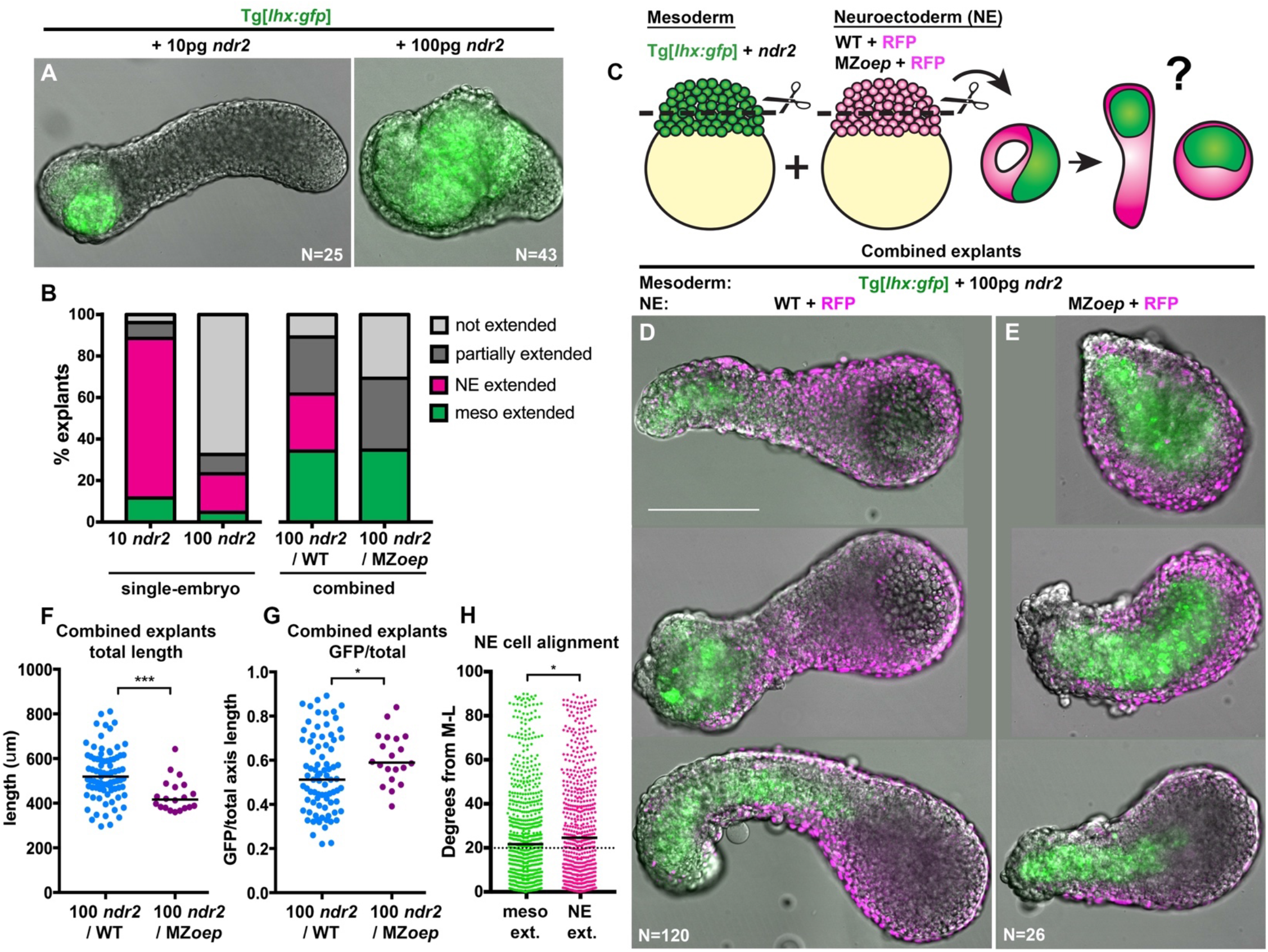
Nodal signaling promotes *ex vivo* neuroectoderm C&E tissue-autonomously. **A**) Representative images of *Tg*[*lhx1a:gfp*] explants injected with 10 or 100pg *ndr2* RNA at the equivalent of 2-4 somite stage. **B**) Mode of extension in single-embryo (left) and combined (right) explants. Number of explants in each condition is indicated in the corresponding panels in (A,D,E). **C**) Diagram of procedure to generate combined explants. **D-E**) Representative images of 100ndr2/WT (D) and *100ndr2/MZ*oep** (E) combined explants. Green is mesoderm labeled by *lhx1a:gfp*. **F-G**) Total length (F) or length of GFP+ region/ total length (G) of combined explants depicted in (D-E). Each dot represents a single explant, black bars are median values, number of explants in each condition is indicated on the corresponding panel in (D-E). ***p=0.0002, *p=0.038, Mann-Whitney test. **H**) Neuroectoderm cell alignment (as in Figure 1) in 100*ndr*2/WT combined explants exhibiting mesoderm (left) or neuroectoderm (right) -driven extension. *p=0.013, K-S test. Scale bar is 200μm.

To determine whether *ndr2*-expressing explants containing only mesoderm extend in the absence of neuroectoderm, we explanted *Tg*[*lhx1a:gfp*] embryos injected with a ten-fold higher dose (100pg) of *ndr2* that produced large amounts of mesoderm and endoderm but very little neuroectoderm (Figure 6A, Figure S3A). This high dose of *ndr2* converted nearly the entire explant into GFP+ mesoderm but yielded far fewer extending explants (~23%) (Figure 6A-B), further supporting our hypothesis that *ndr2*-induced *ex vivo* extension is largely neuroectoderm-driven. However, high doses of Nodal produce mesoderm that differs not only quantitatively (more mesoderm), but also qualitatively (different mesoderm) from that induced by lower doses. For example, 100 pg *ndr2* induced organizer tissue which exhibits little C&E *in vivo* (Ulrich et al., 2003) and expresses *gsc* (Figure S3A), which was shown to inhibit C&E (Ulmer et al., 2017). This raises the possibility that the lack of neuroectoderm alone my not account for the failure of these explants to extend. However, because even extension-prone mesoderm types induced by 10pg *ndr2* often fail to extend *ex vivo* (Figure 6A-B), these results are consistent with *ndr2* driving *ex vivo* extension largely through C&E of neuroectoderm.

### ex vivo neuroectoderm C&E requires tissue-autonomous Nodal signaling

Although mesoderm within *ndr2*-expressing explants does not often itself extend, it is possible that it produces secondary downstream signals that promote C&E of the adjacent neuroectoderm. To test whether Nodal signaling is required tissue-autonomously within the neuroectoderm for its extension, we constructed combined explants in which future ‘mesodermal portions’ consisting of *Tg*[*lhx1a:gfp*] blastoderm expressing high doses (100pg) of *ndr2* were combined with WT or MZ*oep*-/- future ‘neuroectoderm portions’ expressing only a fluorescent marker (RFP) (Figure 6C). As observed in single-origin explants expressing 10pg *ndr2*, many *ndr2/WT* combined explants (~27%) exhibited a mass of GFP+ mesoderm at the end of an extended stalk of RFP+ cells (Figure 6B,D), which were confirmed to express the neuroectoderm marker *sox2* by WISH (Figure S6.1). Critically, this contribution of neuroectoderm to *ex vivo* extension was Nodal-dependent, as *ndr2*/MZ*oep* combined explants never exhibited this configuration of extended neuroectoderm adjacent to an un-extended mass of mesoderm (Figure 6B,E). Furthermore, the total length of combined explants was significantly decreased in *ndr2*/MZ*oep* combined explants compared with *ndr2*/WT (p=0.0002, Mann-Whitney test), while GFP+ mesoderm comprised a larger portion of their axial length (p=0.038, M-W test)(Figure 6F,G), demonstrating that GFP-neuroectoderm contributed little to extension of *ndr2*/MZ*oep* combined explants. Surprisingly, an increased proportion (~34%) of combined explants exhibited robust extension of GFP+ mesoderm (Figure 6B), and this increase was similar regardless of whether the neuroectoderm portion was WT or MZ*oep*-/- (Figure 6B). Together these results demonstrate that *ex vivo* extension of zebrafish tissues can occur by one of two distinct modes of C&E: mesoderm- and/or neuroectoderm-driven. Moreover, neuroectoderm extension requires reception of Nodal signaling in a cell-autonomous manner.

Because the mesoderm portions of combined explants expressed a high dose of *ndr2* that rarely promoted mesoderm extension in single-embryo explants (Figure 6A-B), the observation of extended mesoderm within combined explants raises the possibility that neuroectoderm may promote C&E of neighboring mesoderm by acting as a ‘sink’ for the ligands produced within the mesoderm portion to promote a gradient of Nodal signaling. To test for this possibility, we stained for pSmad2 in 8 hpf 100 *ndr2*/WT combined explants and single-embryo explants injected with 10 or 100pg *ndr2*. Consistent with their robust extension and the results shown in Figure 4, 10pg *ndr2* explants exhibited spatial asymmetry of pSmad2+ nuclei and a gradient of pSmad2 staining intensity (Figure S6.2). Explants injected with 100pg *ndr2* exhibited increased pSmad2 staining intensity and reduced spatial asymmetry of pSmad2+ nuclei compared with 10pg explants, but staining intensity was still graded across the explant, with a slope very similar to that induced by 10pg *ndr2* (Figure S6.2). This indicates that although Nodal activity was detected throughout these explants, the level of signaling activity was still graded. As expected, combining these 100pg *ndr2* explants with an uninjected portion resulted in increased spatial asymmetry of pSmad2+ nuclei and reduced overall staining intensity (Figure S6.2). Surprisingly, however, this did not coincide with a steeper pSmad2 intensity gradient compared to single-embryo 10 or 100pg *ndr2* explants, as the slope of this correlation was similar in all three conditions (Figure S6.2). These results suggest that juxtaposition with uninjected WT blastoderm does not promote mesoderm extension by steepening the Nodal signaling gradient, but that it may do so by reducing overall Nodal levels within the entire explant, or through as yet unexplored mesoderm-neuroectoderm interactions.

We hypothesized that in instances of mesoderm-driven *ex vivo* C&E described above, the overlying neuroectoderm may extend passively along with the underlying mesoderm, which we would expect to stretch neuroectoderm cells along the axis of extension (AP) rather than aligning orthogonal to it (ML). To test this, we measured cell polarity within the neuroectoderm of combined WT explants exhibiting either mesoderm- or neuroectoderm-driven extension. Contrary to our expectations, neuroectoderm cells were actually more ML aligned in cases of mesoderm-driven C&E (Figure 6H), implying that neuroectoderm cells actively polarize regardless of which tissue drives extension. Together with our observation that combined explants containing Nodal-deficient neuroectoderm are significantly less extended than those with WT neuroectoderm (Figure 6F), this supports the notion that Nodal-dependent C&E of the neuroectoderm actively contributes to *ex vivo* axis extension in both the presence and absence of mesoderm-driven C&E.

### Nodal signaling promotes C&E in the absence of mesoderm

The results above demonstrate that the neuroectoderm contributes actively to *ex vivo* axis extension, for which Nodal signaling is required tissue-autonomously. However, because these explants contain a mesodermal component, it remains possible that mesoderm is required for C&E of the adjacent neuroectoderm. We therefore sought to create explants capable of receiving Nodal signals but lacking mesoderm by treating them with a time course of the Nodal inhibitor SB505124 (SB) (Figure 7A). Addition of 50μM SB to *ndr2*-expressing explants at 4, 5, or 6 hpf completely blocked both explant extension (Figure 7A-B) and expression of the mesoderm markers (and direct transcriptional targets of Nodal signaling, (Dubrulle et al., 2015)) *tbxta, noto*, and *tbx16* at 2-4S stage (Figure 7C). Because mesoderm marker expression was detectable by WISH in untreated *ndr2*-injected explants beginning at 5 hpf (Figure S7.1), these results imply that Nodal signaling is required for both specification and maintenance of mesodermal fates *ex vivo*. Explants treated with SB at 8 hpf (concurrent with C&E onset) underwent extension but were significantly shorter than DMSO-treated controls despite robust mesoderm marker expression (Figure 7B-C) (p<0.0001, Mann-Whitney test). Notably, this differed from SB treatment of intact WT embryos, which did not disrupt mesoderm marker expression or AP axis extension after 5hpf (Figure S7.2)(Hagos & Dougan, 2007). Because loss of Nodal signaling prevented full explant extension even in the presence of mesoderm, these results further support a role for Nodal in *ex vivo* C&E, distinct from its role in mesoderm formation.

**Figure 7:**
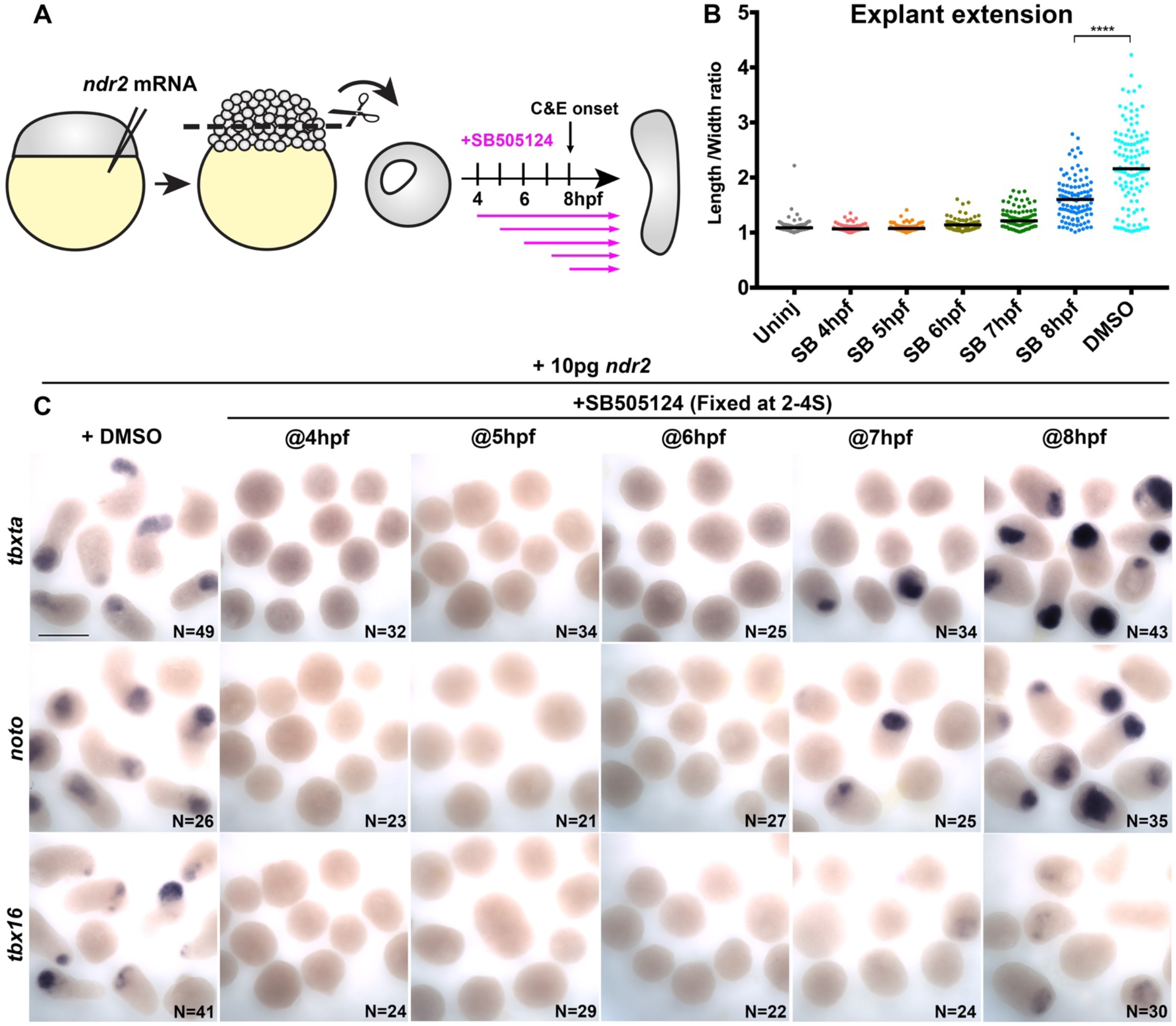
Nodal inhibition after mesoderm formation prevents full explant extension. **A**) Diagram of time course of SB505124 (SB) treatment of *ndr2*-injected explants. **B**) Length/width ratios of WT explants injected with 10pg *ndr2* RNA, treated with SB at the indicated time points, and fixed at the equivalent of 2-4 somite stage (depicted in (C)). Each dot represents a single explant, black bars are median values, p<0.0001, Mann-Whitney test. **C**) In situ hybridization for the transcripts indicated in explants described in (B). Scale bar is 300μm.

This experiment has not, however, allowed us to assess Nodal-dependent neuroectoderm extension in the absence of mesoderm. We therefore treated *ndr2*-expressing explants with SB in discrete two-hour windows followed by wash-out of the inhibitor (Figure 8A). Treatment from 4-6 hpf completely blocked mesoderm marker expression and extension, even after the drug was removed (Figure 8B-C). Treatment from 6-8 hpf similarly blocked expression of mesoderm markers, but these explants exhibited marked (albeit reduced) extension (Figure 8B-C). Because sustained SB treatment beginning at 6 hpf completely blocked explant extension (Figure 7), this extension must be driven by Nodal signaling after removal of the inhibitor at 8 hpf. Indeed, inhibiting only this later phase of Nodal signaling by SB treatment at 8hpf prevented full explant extension (Figures 7B, 8B), confirming that Nodal signaling after C&E onset contributes significantly to *ex vivo* extension morphogenesis.

**Figure 8:**
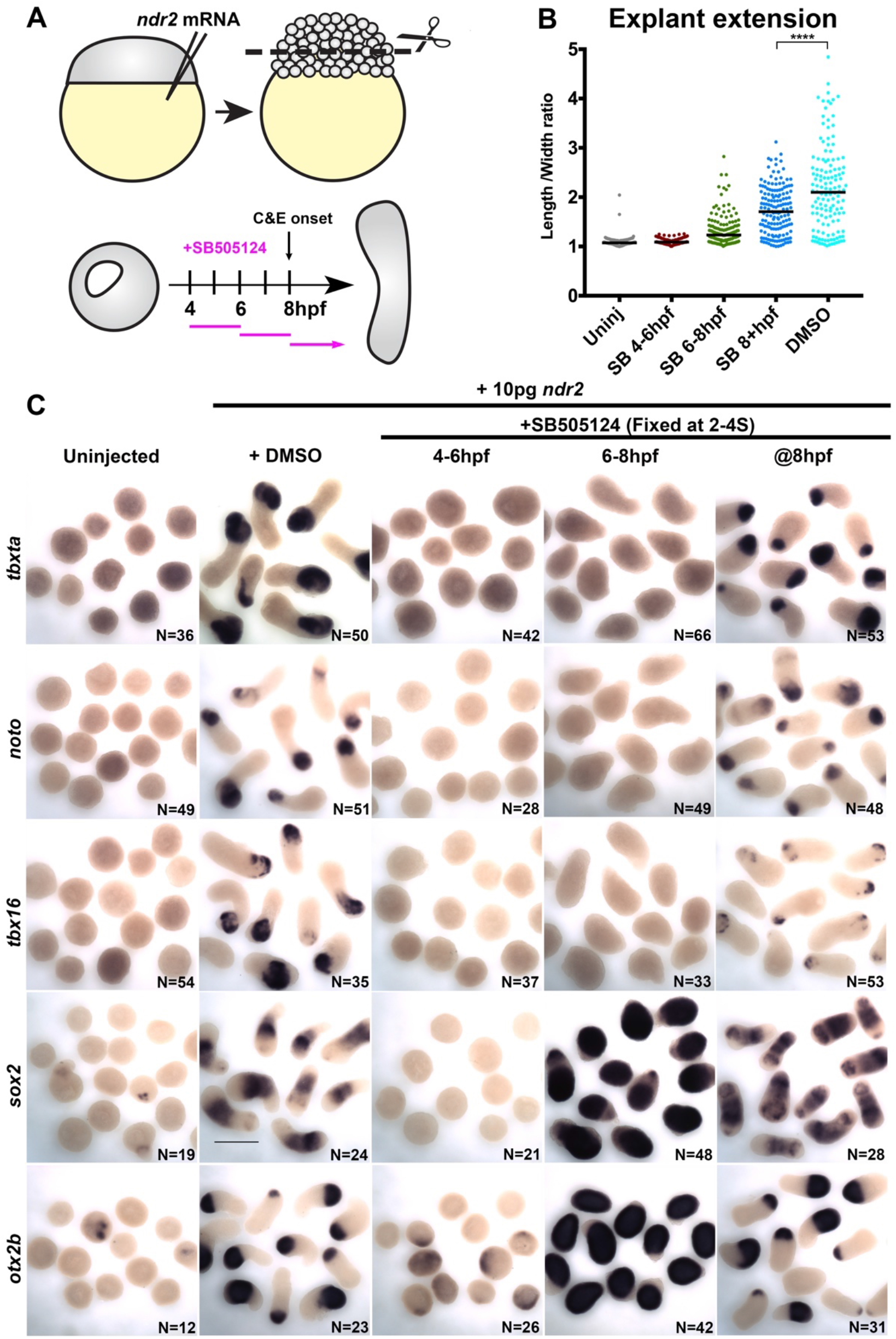
Nodal promotes *ex vivo* neuroectoderm C&E in the absence of mesoderm. **A**) Diagram of time course of SB treatment of *ndr2*-injected explants. **B**) Length/width ratios of WT explants injected with 10pg *ndr2* RNA, treated with SB at the indicated time points, and fixed at the equivalent of 2-4 somite stage (depicted in (C), as in Figure 7). p<0.0001, Mann-Whitney test. **C**) In situ hybridization for the transcripts indicated in explants described in (B). Scale bar is 300μm.

Because explants treated with SB from 6-8 hpf exhibited Nodal-dependent extension but did not express mesoderm markers, we hypothesized that the observed extension was driven by neuroectoderm. Indeed, we found that explants treated with SB from 6-8 hpf exhibited dramatically increased expression of neuroectoderm markers *sox2* and *otx2b* by WISH compared with DMSO-treated controls (Figure 8C), demonstrating that Nodal signaling promotes C&E of neuroectoderm even in the absence of mesoderm. Expression of neuroectoderm markers was also detected in explants treated at 8hpf but was dramatically reduced or absent in those treated at 4-6 hpf, consistent with the previous finding that unmanipulated zebrafish blastoderm explants differentiate into enveloping layer rather than neuroectoderm (Sagerström, Gammill, Veale, & Sive, 2005). Together these results provide evidence for a sustained requirement for Nodal signaling during *ex vivo* axis extension, including a late phase after C&E onset that promotes mesoderm-independent extension.

## DISCUSSION

Coordination of embryonic patterning and morphogenesis is among the most fundamental and least understood questions in developmental biology. Convergence and extension (C&E) are evolutionarily conserved gastrulation movements that narrow germ layers mediolaterally and elongate them along the AP axis. AP axial patterning is necessary for C&E (Ninomiya et al., 2004), but how C&E movements are coordinated with embryonic patterning is only beginning to be understood. The noncanonical Wnt/PCP signaling network is thought to act as a molecular compass to mediate ML cell polarization (Gray et al., 2011). Links between PCP and morphogens that pattern the embryo have been identified in zebrafish gastrulae, including the ventral to dorsal gradient of BMP signaling that both specifies ventral cell fates (Nguyen et al., 1998) and prevents C&E movements in ventral regions of the gastrula in part by limiting expression of Wnt/PCP components (Myers, Sepich, & Solnica-Krezel, 2002) and regulating cell adhesion (von der Hardt et al., 2007). However, signals that orient the PCP compass to promote polarized C&E cell behaviors remain poorly defined. Here, using a combination of intact zebrafish gastrulae and an *ex vivo* model of axial extension, we have characterized a critical role for the morphogen Nodal in regulation of cell behaviors underlying embryonic axis extension. Although previous studies found that Nodal regulates neuroectoderm morphogenesis indirectly by specifying mesoderm (Aquilina-Beck et al., 2007; Araya et al., 2014; Gonsar et al., 2016), we have identified an additional mesoderm-independent role for Nodal signaling in promoting cell polarity and C&E morphogenesis within the neuroectoderm. Moreover, we find that Nodal-dependent neuroectoderm C&E contributes significantly to a simplified model of axis extension, demonstrating functional importance of both mesoderm-dependent and -independent Nodal function.

We provide several lines of evidence to support this role for Nodal signaling in tissue extension independent of mesoderm. First, ML polarization of Nodal-deficient neuroectoderm cells is not completely restored upon transplantation into WT hosts, demonstrating a cell-autonomous function (Figure 1). Second, exogenous Nodal ligands promote neuroectoderm extension in otherwise naïve blastoderm explants (Figure 6). Third, Nodal signaling is required tissue-autonomously for this neuroectoderm extension (Figure 6). Finally, Nodal ligands are required for full explant extension even in the presence of mesoderm, and promote partial extension in the absence of mesoderm (Figures 7-8). Additional evidence demonstrates that this Nodal-dependent neuroectoderm C&E actively contributes to *ex vivo* axis extension. First, although combined explants containing WT mesoderm and Nodal-deficient neuroectoderm can extend by C&E of the mesoderm alone, they are significantly shorter than explants in which both the mesoderm and neuroectoderm extend (Figure 6). Then, even in cases of mesoderm extension, WT neuroectoderm cells become ML polarized, suggesting that they are actively intercalating. Our results also highlight the important role that mesoderm plays in axis extension, however. For example, mesoderm in WT host embryos can partially, although not fully, restore ML polarity to transplanted Nodal-deficient neuroectoderm cells (Figure 1), and explants containing both mesoderm and neuroectoderm extend better than those containing neuroectoderm alone (Figure 8). The important role of vertical interactions between mesoderm and neuroectoderm in C&E has long been appreciated ((Elul & Keller, 2000; Elul, Koehl, & Keller, 1997; Ezin, Skoglund, & Keller, 2006), and we will be interested to examine how such interactions influence morphogenesis *ex vivo*, and whether this differs depending on the configuration of mesoderm (either vertically or in plane with the neuroectoderm) within our explants.

Previous work found that asymmetric Activin signaling is sufficient to promote C&E *ex vivo* (Ninomiya et al., 2004). We found that closely-related Nodal ligands are similarly sufficient for *ex vivo* extension, and furthermore, that Nodal pathway activation was asymmetric within our *ndr2*-expressing explants (Figure 4). At least two non-mutually exclusive hypotheses could explain how asymmetric Nodal signaling is translated into axis extension. First, Nodal could activate tissue-specific gene expression networks that both define tissue type and include instructions for C&E cell behaviors that are specific to each tissue. Alternatively, Nodal could promote differential expression of instructive signaling molecules along the axis of extension independent of the tissue types specified. This second possibility is supported by evidence that Nodal pathway ligands can induce extension at a number of different doses, irrespective of the tissue types produced (Figure S3), so long as that signaling is asymmetric (Ninomiya et al., 2004). Importantly, although we observe asymmetric Nodal signaling in extending explants, it is not yet clear that this asymmetry is required for extension.

As discussed above, non-canonical Wnt/PCP signaling is thought to act as a molecular compass that orients cells with respect to the embryonic axes (Gray et al., 2011). Because Nodal signaling is necessary for both AP axial patterning (Chen & Schier, 2001; Feldman et al., 2000; B. Thisse et al., 2000) and C&E cell behaviors (this work), it is a prime candidate to act upstream of PCP to orient this compass, thereby coordinating axial patterning and morphogenesis. Indeed, like others (Ninomiya et al., 2004), we found that Nodal-induced *ex vivo* C&E is reduced when PCP signaling is disrupted (Figure 5), suggesting that Nodal may regulate C&E through downstream PCP signaling. However, additional *in vivo* evidence suggests a more complicated interaction between these two signaling pathways. First, we found that transcripts encoding PCP signaling components are expressed normally within Nodal-deficient MZ*oep* mutant gastrulae (Figure 2), demonstrating that Nodal is unlikely to influence PCP via transcriptional regulation. We further found that localization of the core PCP component Prickle is only mildly affected in MZ*oep* mutants (Figure 2), suggesting that although Nodal is required for polarization of cell behaviors (Figure 1), it is not required for asymmetric distribution of this core PCP component. Finally, axis extension defects resulting from complete loss of Nodal signaling are exacerbated by disrupted PCP signaling (Figure 3), implying that PCP signaling retains some residual polarizing behavior in the absence of Nodal, just as Nodal retains some polarizing activity upon disruption of PCP (Figure 5). Taken together, these data demonstrate that PCP and Nodal each provide some function or cue that is necessary - but not sufficient - for full ML polarization of cells and cell behaviors underlying C&E, suggesting that they operate via partially parallel but cooperating pathways. This is also consistent with our transplant results, in which cell-autonomous Nodal signaling was required for full planar polarization at the end of gastrulation, but an additional Nodal-independent mechanism was still functional during an earlier phase (Figure 1). Notably, results from our explants lead us to different conclusions regarding the relationship between Nodal and PCP. Within explants, Nodal ligands are sufficient to promote PCP-dependent cell polarization and C&E in the absence of other patterning cues (Figures 3,5), indicating that PCP functions wholly downstream of Nodal signaling in this *ex vivo* context. C&E and cell polarization are not completely abolished in *ndr2*-expressing *vangl2* morphant explants, however, demonstrating that Nodal likely contributes to polarized cell behaviors through an additional PCP-independent mechanism.

The Nodal-expressing explants described in this study provide a robust, simplified model of axial extension in which a signaling molecule of interest – in this case, Nodal – can be studied independently of endogenous patterning and signaling events. Although this is a powerful tool, it is important to acknowledge the ways in which this system differs from intact embryos. First, we found that the period of detectable Nodal signaling activity was substantially earlier and shorter in embryos than explants, as observed by pSmad2 immuno-staining and responses to SB treatment. Nodal inhibition through 7 hpf effectively blocked extension in explants, whereas extension of intact embryos was only reduced when treated at 5 hpf or earlier, consistent with longer and later Nodal signaling in explants. Second, because explants do not contain the full complement of molecular signals and tissue interactions present in intact embryos, the contribution of additional signals to C&E may be masked by the reliance of explant extension on Nodal alone. This is illustrated, for example, by the function of PCP in parallel with Nodal *in vivo* but strictly downstream of Nodal *ex vivo*. The absence of a yolk cell also dramatically alters the geometry of explanted tissues and removes signaling input from the extraembryonic yolk syncytial layer. Despite these differences, explants exhibit a suite of complex, biologically relevant behaviors in common with intact embryos, including C&E morphogenetic movements, ML cell polarization, timing of C&E onset, and transcriptional responses to Nodal. These explants, which we have dubbed “morphonoids”, therefore provide a simplified, synthetic platform that has allowed for new insights into the role of Nodal signaling in C&E morphogenesis.

## MATERIALS & METHODS

### Zebrafish

Adult zebrafish were raised and maintained according to established methods (Westerfield, 1993) in compliance with standards established by the Washington University Animal Care and Use Committee. Embryos were obtained from natural mating and staged according to morphology as described (Kimmel, Ballard, Kimmel, Ullmann, & Schilling, 1995). All studies on WT were carried out in AB* backgrounds.

Additional lines used include *oep^tz257^* (Hammerschmidt et al., 1996) and *Tg*[*lhx1a:gfp*] (Swanhart et al., 2010). *oep-/-* embryos were rescued by injection of 50pg synthetic *oep* RNA and raised to adulthood, then intercrossed to generate maternal-zygotic *oep-/-* embryos. Fish were chosen from their home tank to be crossed at random, and the resulting embryos were also chosen from the dish at random for injection and inclusion in experiments.

### Microinjection of synthetic RNA and morpholino oligonucleotides

One-celled embryos were aligned within agarose troughs generated using custom-made plastic molds and injected with 1-3 pL volumes using pulled glass needles. Synthetic mRNAs for injection were made by *in vitro* transcription from linearized plasmid DNA templates using Invitrogen mMessage mMachine kits. Doses of RNA per embryo were as follows: 100pg *membrane Cherry*, 50pg membrane *eGFP*, 25pg *H2B-RFP*, 50pg *pk-gfp*, and 2.5-100pg *ndr2*. Injection of 2ng *MO1-tri/vangl2* (Williams et al., 2012) was carried out as for synthetic RNA.

### Immunofluorescent staining

Embryos were stained for phosphorylated Smad2 as described in (van Boxtel et al., 2015). Briefly: embryos and explants were fixed overnight in 4% PFA, rinsed in PBT, and dehydrated to 100% methanol. Prior to staining, embryos were rehydrated into PBS, rinsed in PBS + 1% Triton X-100, and incubated in ice-cold acetone at -20°C for 20 minutes. Embryos/explants were then blocked in PBS+ 10% FBS and 1% Triton X-100 and incubated overnight at 4°C with an anti-pSmad2/3 antibody (Cell Signaling #8828) at 1:1000 in block. Samples were rinsed in PBT/1% Triton X-100 and incubated with Alexa Fluor 488 anti-Rabbit IgG (Invitrogen) at 1:1000. Embryos were co-stained with 4’,6-Diamidino-2-Phenylindole, Dihydrochloride (DAPI) and rinsed in PBS/1% Triton X-100 prior to mounting in 2% methylcellulose for confocal imaging.

### Whole mount in situ hybridization

Antisense riboprobes were transcribed using NEB T7 or T3 RNA polymerase and labeled with digoxygenin (DIG) (Roche). Whole-mount *in situ* hybridization (WISH) was performed according to (C. Thisse & Thisse, 2008). Briefly: Embryos were fixed overnight in 4% paraformaldehyde (PFA) in phosphate buffered saline (PBS), rinsed in PBS + 0.1% tween 20 (PBT), and dehydrated into methanol. Embryos were then rehydrated into PBT, incubated for at least two hours in hybridization solution with 50% formamide (in 0.75 M Sodium chloride, 75mM Sodium citrate, 0.1% tween 20, 50μg/mL Heparin (Sigma), and 200μg/mL tRNA) at 70°C, then hybridized overnight at 70°C with antisense probes diluted approximately 1ng/μl in hybridization solution. Embryos were washed gradually into 2X SSC buffer (0.3 M Sodium chloride, 30mM Sodium citrate), and then gradually from SSC to PBT. Embryos were blocked at room temperature for several hours in PBT with 2% goat serum and 2 mg/mL bovine serum albumin (BSA), then incubated overnight at 4°C with anti-DIG antibody (Roche #11093274910) at 1:5000 in block. Embryos were rinsed extensively in PBT, and then in staining buffer (PBT +100mM Tris pH 9.5, 50mM MgCl2, and 100mM NaCl) prior to staining with BM Purple solution (Roche).

### Blastoderm explants

Embryos were injected with *ndr2* RNA (and MOs) at one-cell stage as described above and dechorionated using Pronase (Roche). At 256- to 512-cell stage, watchmaker’s forceps were used to excise the animal portion of each embryo in an agarose-coated dish filled with 3X Danieau solution. Explants were allowed to heal briefly, then transferred into agarose-coated 6-well plates containing explant medium - comprised of Dulbecco’s modified eagle medium with nutrient mixture F-12 (Gibco DMEM/F12) containing 2.5mM L-Glutamine, 15mM HEPES, 3% newborn calf serum, and penicillin-streptomycin - and incubated at 28.5°C. For combined explants, the newly-excised blastoderm of two embryos were placed cut-side together and allowed to heal briefly in 3X Danieau before transferring to culture medium.

### Pharmacological treatments

50μM SB505124 (Sigma #S4696) was added to the media of explants (and embryos) in agarose-coated 6-well plates at the times specified. For wash-out experiments, SB-containing medium was removed and explants were washed twice with 0.3x Danieau solution before replacement with fresh explant medium.

### Transplantation

For cell autonomy and Pk-GFP transplants, host and donor embryos were injected with RNA encoding different fluorescent markers and/or Pk-GFP as described above and dechorionated using Pronase. Host and donor embryos were arranged within the wells of a custom-molded agarose plate at approximately sphere stage, and approximately 20-40 cells were transferred from donors to hosts using a fine-pulled glass capillary.

### Microscopy

Live embryos/explants expressing fluorescent proteins were mounted in 0.75% low-melt agarose, and fixed embryos/explants subjected to immunofluorescent staining were mounted in 2% methylcellulose in glass-bottomed 35-mm petri dishes for imaging using a modified Olympus IX81 inverted spinning disc confocal microscope equipped with Voltran and Cobolt steady-state lasers and a Hamamatsu ImagEM EM CCD digital camera. For live time-lapse series, 60 μm z-stacks with a 2μm step were collected every three to five minutes (depending on the experiment) for three hours using a 20 x or 40x dry objective lens for intact embryos and a 20x objective for explants. Temperature was maintained at 28.5°C during imaging using a Live Cell Instrument stage heater. For immunostained embryos and explants, 100 μm z-stacks with a 1 or 2μm step were collected using a 10x or 20x dry objective lens, depending on the experiment. Bright field and transmitted light images of live embryos and *in situ* hybridizations were collected using a Nikon AZ100 macroscope.

### Image analysis

ImageJ was used to visualize and manipulate all microscopy data sets. For cell shape measurements in live embryos, a single z-plane (in ubiquitously labeled embryos) or a projection of several z-planes (when measuring transplanted cells) through the neuroectoderm was chosen for each time point. To measure cell orientation and elongation, the anteroposterior axis in all embryo images was aligned prior to manual outlining of cells. A fit ellipse was used to measure orientation of each cell’s major axis and its aspect ratio. To assess Pk-GFP localization, isolated donor cells expressing Pk-GFP were scored according to subcellular localization of GFP signal. For cell tracking, Imaris software and the ImageJ TrackMate plugin were used to automatically detect and track labeled nuclei in the dorsal hemisphere of WT and MZ*oep* mutant gastrulae and to produce color-coded depictions of their trajectories. To measure spatial distribution of pSmad2 immunostaining in explants, maximum intensity z-projections were made from each confocal stack for both pSmad2 and DAPI channels. All images were oriented such that the highest apparent pSmad2 signal (if any) was to the left, and the ImageJ ‘Colocalization’ plugin was used to detect colocalization of pSmad2 and DAPI channels. Next, the TrackMate plugin was used to detect both colocalized and DAPI-alone spots, whose X locations were quantified. To measure length/width ratios of explants, we divided the length of a segmented line drawn along the midline of each explant (accounting for curvature) by the length of a perpendicular line spanning the width of the explant near its midpoint. To measure width of the neural plate in whole mount embryos, dorsal-view images were collected of each embryo, and a line was drawn from one side of the *dlx3b* expression domain to the other side at the level of the mid-hindbrain boundary marked by *fgf8* expression. Length measurements were made similarly by measuring from the anterior to posterior aspects of the *dlx3b* expression domain in lateral-view images. Images were coded and analyses were performed blinded to ensure unbiased measurements.

### Statistical analysis

Graphpad Prism 7 software was used to perform statistical analyses and generate graphs of data collected from embryo and explant images. The statistical tests used varied as appropriate for each experiment and are described in the text and figure legends. Data were tested for normal distribution, and non-parametric tests (Mann-Whitney and Kolmogorov-Smirnov) were used for all non-normally distributed data. Normally distributed data with similar variance between groups were analyzed using parametric tests (T-tests and ANOVAs). All tests used were two-tailed.

## ACKNOWLEDGEMENTS

We thank Dr. Alex Schier and lab members for generously sharing fish and reagents, and Drs. Diane Sepich, Ann Sutherland, Ray Keller, and Dave Shook for helpful discussions. This work was supported by National Institutes of Health awards K99HD091386 to MLKW and 1R35GM118179 to LSK.

