## Supplemental figures for "A mesoderm-independent role for Nodal signaling in convergence & extension gastrulation movements"

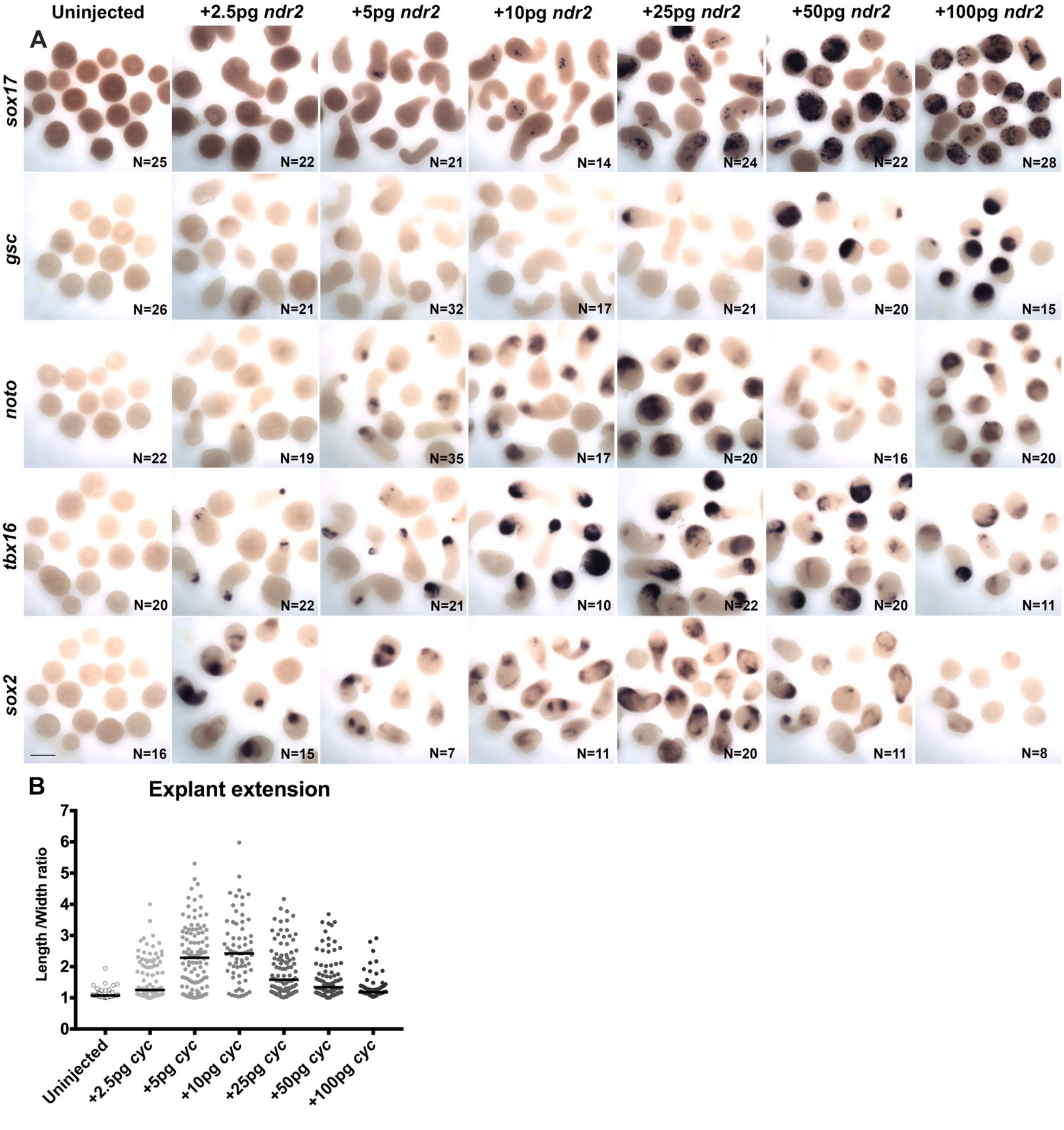


**Figure S3: Nodal ligand levels regulate cell fate and extension of explants.**
**A**) In situ hybridization for the transcripts indicated in WT explants injected with 0-100 pg *ndr2* RNA and fixed at the equivalent of 2-4 somite stage. **B**) Length/width ratios of explants depicted in (A). Each dot represents a single explant, black bars are median values. Scale bar is 200μm.

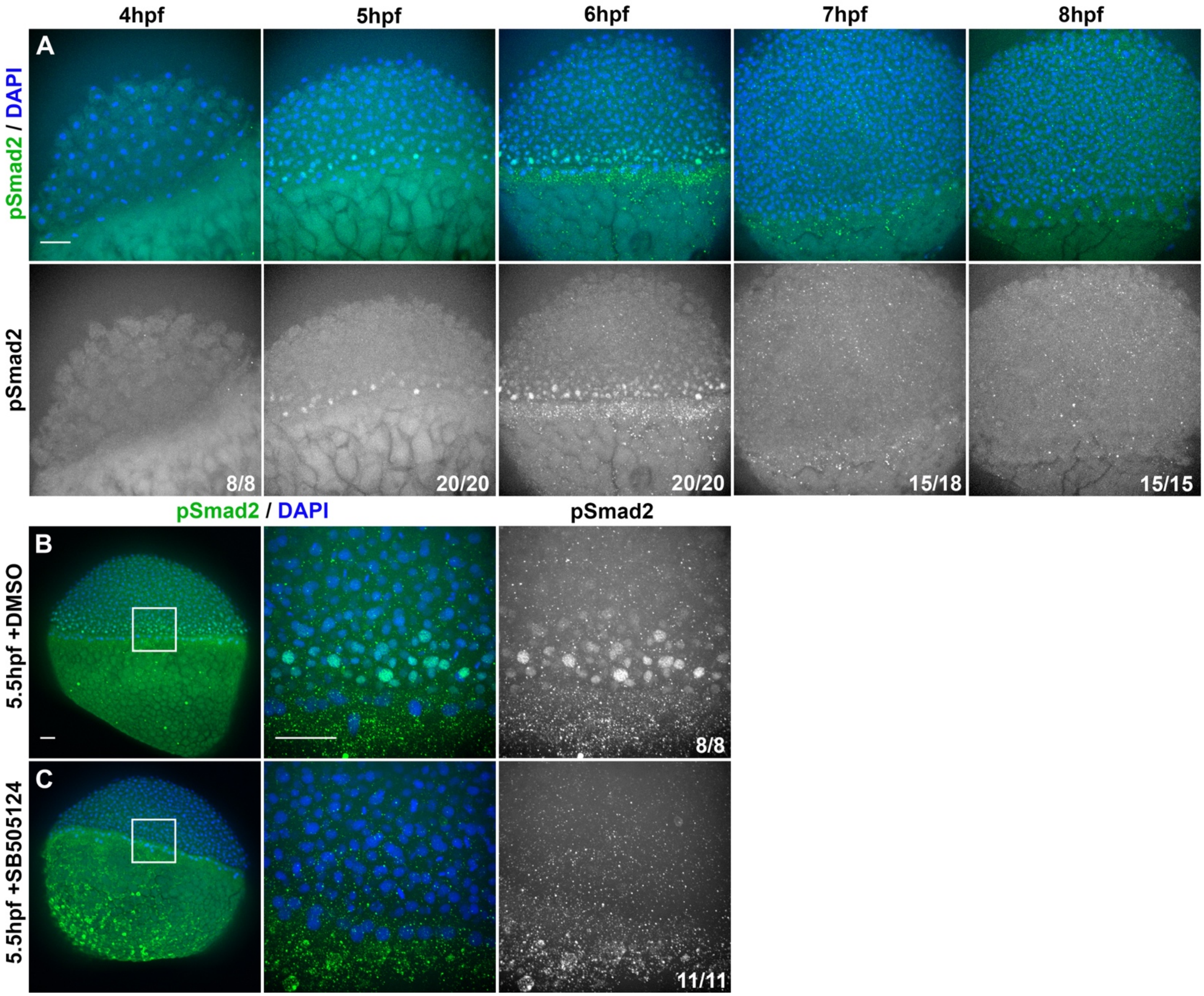


**Figure S4: Nodal signaling activity in intact embryos.**
**A**) Representative confocal z-stacks of immunofluorescent (IF) staining for phosphorylated Smad2 (bottom) overlaid with DAPI-labeled nuclei (top) in WT embryos at the time points indicated. **B**) Representative confocal z-stacks of pSmad2 IF and DAPI staining in 5.5 hpf WT embryos treated with DMSO (top) or SB505124 (bottom) starting at 3 hpf. Panels to the right are magnified views of regions within white squares. Fractions indicate the number of embryos with the depicted phenotype over the total number of embryos examined for each condition. The animal pole is up in all images. Scale bars are 50μm.

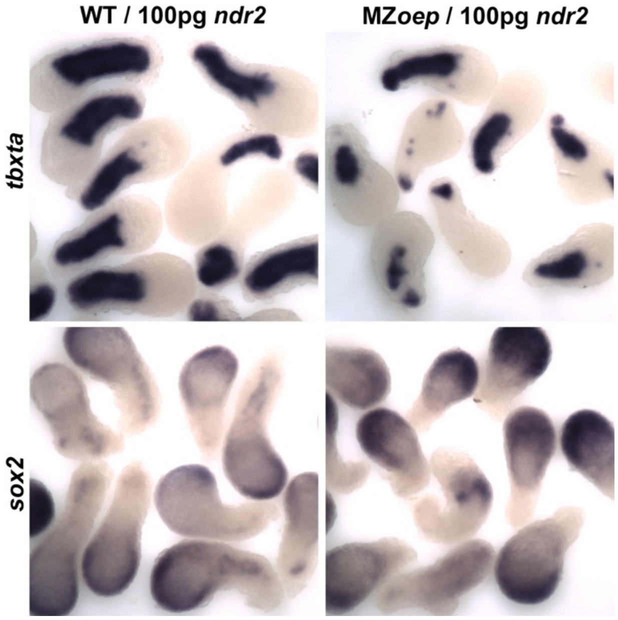


**Figure S6.1: Combined explants contain mesoderm and neuroectoderm.**
In situ hybridization for the mesoderm marker *tbxta* and the neuroectoderm marker *sox2* in combined explants of the genotypes indicated at the equivalent of 2-4 somite stage.

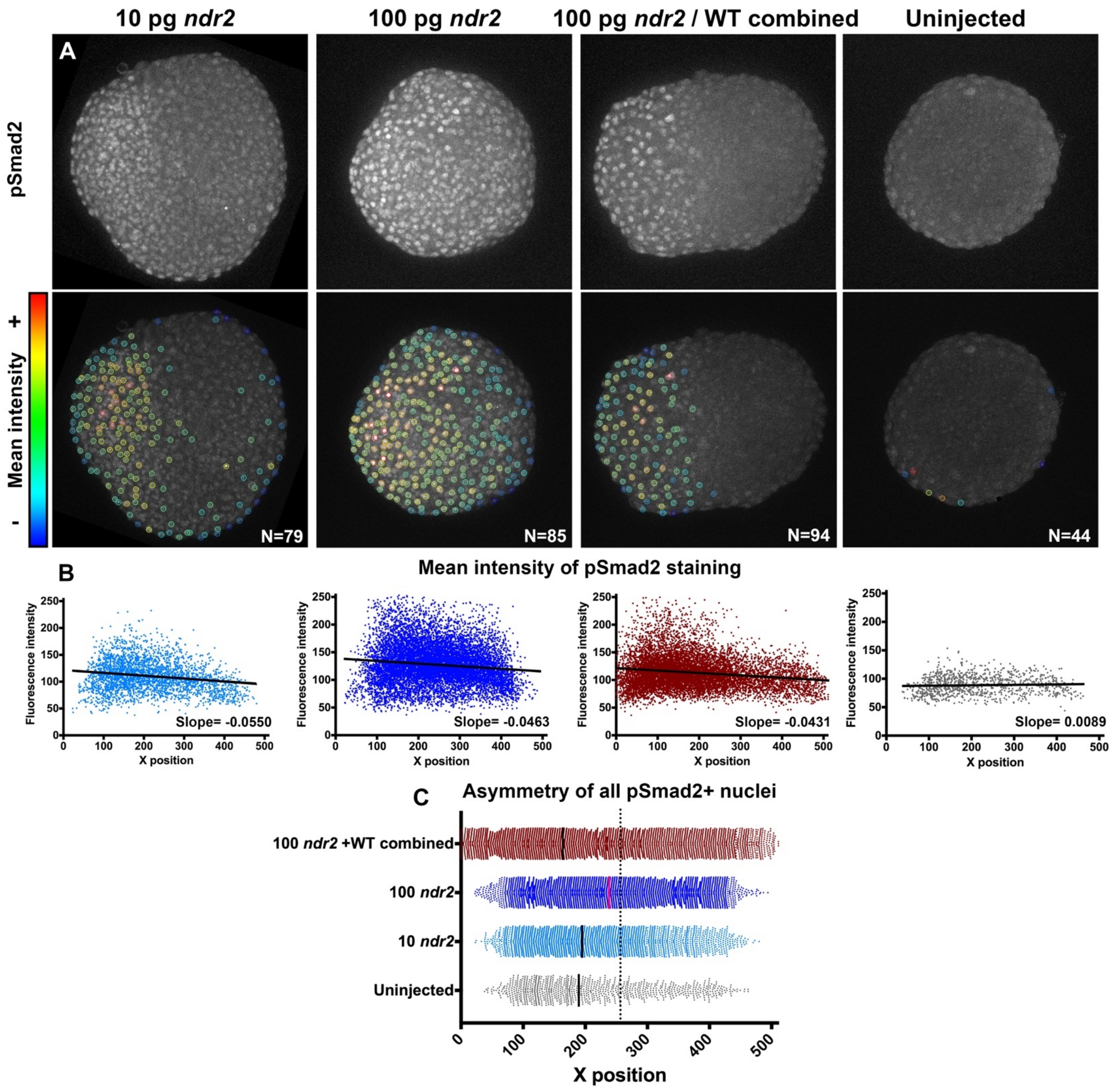


**Figure S6.2: Asymmetry of Nodal signaling activity in single-embryo and combined explants.**
**A**) Representative confocal z-projections of immunofluorescent staining for phosphorylated Smad2 in 8 hpf explants of the conditions indicated. Colored circles indicate automatically detected spots and are colored according to pSmad2 mean staining intensity. **B**) Correlation between mean pSmad2 staining intensity and position within the explant. Black lines are linear regressions with slopes indicated at the bottom of each panel. **C**) Axis position of pSmad2-positive nuclei in 8 hpf explants of the conditions indicated. Bars are median values.

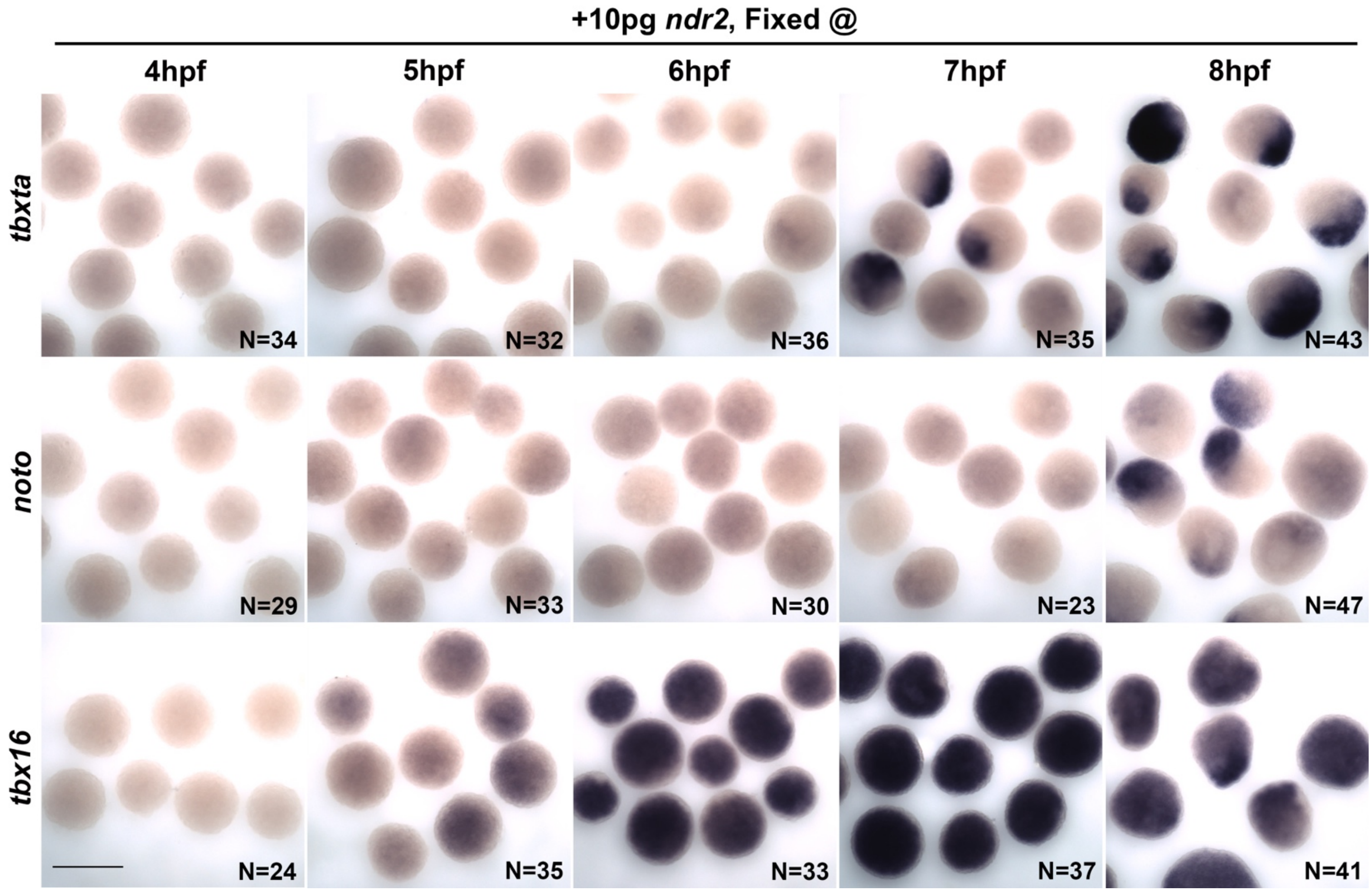


**Figure S7.1: Time course of mesoderm marker expression in explants.**
In situ hybridization for the transcripts indicated in WT explants injected with 10pg *ndr2* RNA and fixed at 4-8 hpf. Scale bar is 300μm.

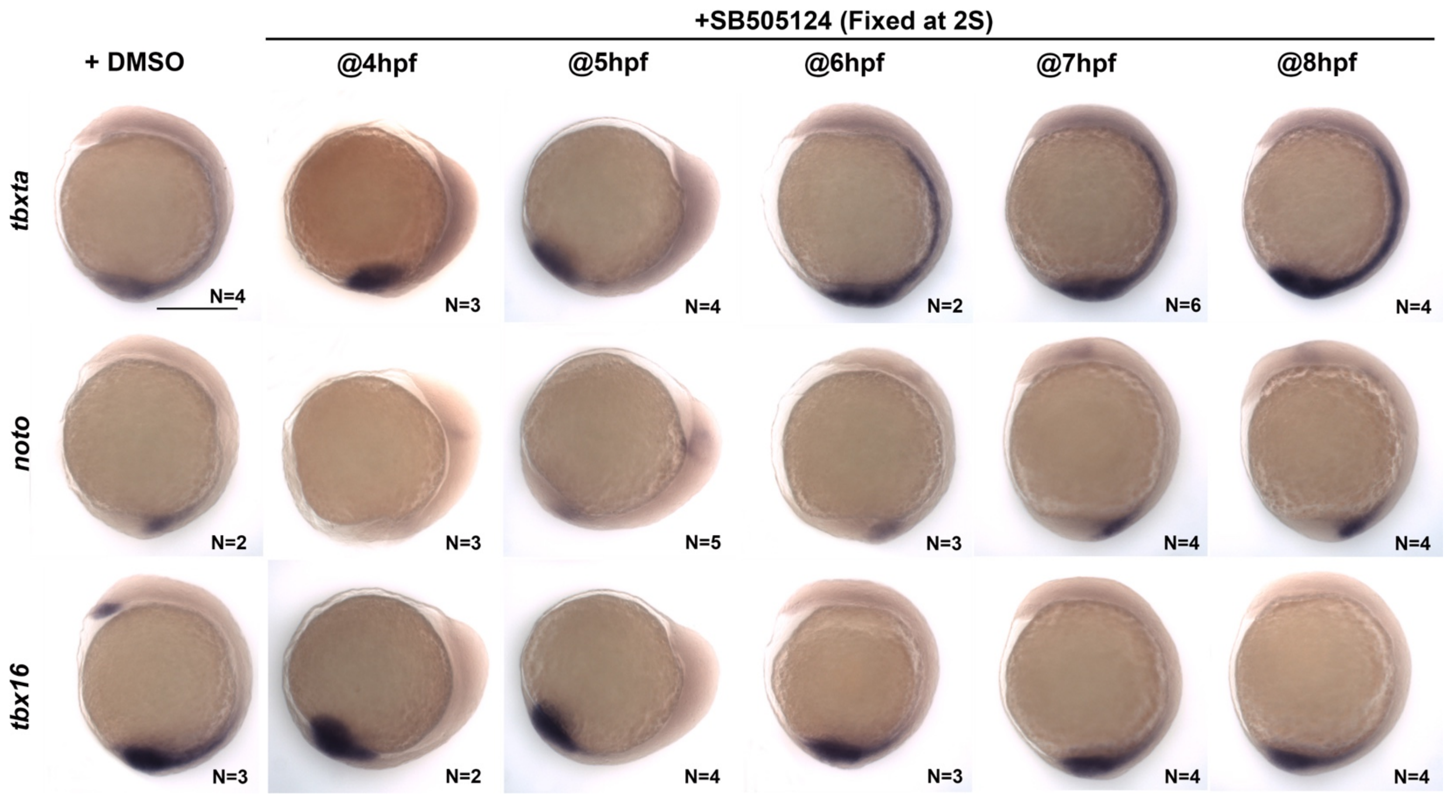


**Figure S7.2: Nodal inhibitor treatment of intact embryos.**
In situ hybridization for the transcripts indicated in WT embryos treated with DMSO or SB505124 beginning at 4-8 hpf and fixed at 2 somite stage. Scale bar is 300μm.
